# Genome-wide association study of Alzheimer’s disease CSF biomarkers in the EMIF-AD Multimodal Biomarker Discovery dataset

**DOI:** 10.1101/774554

**Authors:** Shengjun Hong, Dmitry Prokopenko, Valerija Dobricic, Fabian Kilpert, Isabelle Bos, Stephanie J. B. Vos, Betty M. Tijms, Ulf Andreasson, Kaj Blennow, Rik Vandenberghe, Isabelle Cleynen, Silvy Gabel, Jolien Schaeverbeke, Philip Scheltens, Charlotte E. Teunissen, Ellis Niemantsverdriet, Sebastiaan Engelborghs, Giovanni Frisoni, Olivier Blin, Jill C. Richardson, Regis Bordet, José Luis Molinuevo, Lorena Rami, Alzheimer’s Disease Neuroimaging Initiative (ADNI), Petronella Kettunen, Anders Wallin, Alberto Lleó, Isabel Sala, Julius Popp, Gwendoline Peyratout, Pablo Martinez-Lage, Mikel Tainta, Richard J. B. Dobson, Cristina Legido-Quigley, Kristel Sleegers, Christine Van Broeckhoven, Mara ten Kate, Frederik Barkhof, Henrik Zetterberg, Simon Lovestone, Johannes Streffer, Michael Wittig, Andre Franke, Rudolph E Tanzi, Pieter Jelle Visser, Lars Bertram

## Abstract

Alzheimer’s disease (AD) is the most prevalent neurodegenerative disorder and the most common form of dementia in the elderly. Susceptibility to AD is considerably determined by genetic factors which hitherto were primarily identified using case-control designs. Elucidating the genetic architecture of additional AD-related phenotypic traits, ideally those linked to the underlying disease process, holds great promise in gaining deeper insights into the genetic basis of AD and in developing better clinical prediction models. To this end, we generated genome-wide single-nucleotide polymorphism (SNP) genotyping data in 931 participants of the European Medical Information Framework Alzheimer’s Disease Multimodal Biomarker Discovery (EMIF-AD MBD) sample to search for novel genetic determinants of AD biomarker variability. Specifically, we performed genome-wide association study (GWAS) analyses on 16 traits, including 14 measures of amyloid-beta (Aβ) and tau-protein species in the cerebrospinal fluid (CSF). In addition to confirming the well-established effects of apolipoprotein E (*APOE*) on diagnostic outcome and phenotypes related to Aβ42, we detected novel potential signals in the zinc finger homeobox 3 (*ZFHX3*) for CSF-Aβ38 and CSF-Aβ40 levels, and confirmed the previously described sex-specific association between SNPs in geminin coiled-coil domain containing (*GMNC*) and CSF-tau. Utilizing the results from independent case-control AD GWAS to construct polygenic risk scores (PRS) revealed that AD risk variants only explain a small fraction of CSF biomarker variability. In conclusion, our study represents a detailed first account of GWAS analyses on CSF-Aβ and -tau related traits in the EMIF-AD MBD dataset. In subsequent work, we will utilize the genomics data generated here in GWAS of other AD-relevant clinical outcomes ascertained in this unique dataset.

## Introduction

Alzheimer’s disease (AD) is a progressive and devastating neurodegenerative disorder, which leads to cognitive decline, loss of autonomy, dementia, and eventually death. Neuropathologically, AD is characterized by the accumulation of extracellular amyloid β (Aβ) peptide deposits (“plaques”) and intracellular hyperphosphorylated tau protein aggregates (“tangles”) in the brain [1], [2]. Using genetic linkage analysis followed by positional cloning led to the discovery of rare mutations in three genes encoding the amyloid-beta precursor protein (*APP*) and presenilins 1 and 2 (*PSEN1*, *PSEN2*) that cause fully penetrant monogenic forms of AD [3]. However, the vast majority of patients likely summer from a polygenic (“sporadic”) form of AD which is driven by numerous genomic variants [4], the identification of which are the main aim of genome-wide association studies (GWAS).

The most strongly and most consistently associated AD risk gene (even prior to GWAS [5]) is *APOE*, which encodes apolipoprotein E, a cholesterol transport protein that has been implicated in numerous amyloid-specific pathways, including amyloid trafficking as well as plaque clearance [2], [6]. In addition to *APOE*, nearly three dozen independent loci have now been reported to be associated with disease risk by GWAS [7]–[9]. Pathophysiologically, the risk genes identified to date appear to predominantly act through modulations of the immune system response, endocytotic mechanisms, cholesterol homeostasis, and APP catabolic processes [7], [8].

Despite these general advances in the field of AD genetics, many key questions still remain to be answered. First, even when analyzed in combination, the currently known AD risk factors explain only a fraction of the phenotypic variance [7], and, accordingly, only have limited applicability as early markers for disease onset and progression [7], [10]. Second, most of the currently reported AD susceptibility genes were identified using classic case-control designs comparing clinically manifest dementia-stage AD vs control individuals, typically lacking data on early-stage impairments (e.g. mild cognitive impairment [MCI]) and clinical follow-up to ascertain progression and eventually conversion to AD. Finally, while some studies have investigated the correlation between genetics and non-genetic biomarkers, this was hitherto typically done as bivariate assessments owing to the lack of a broad spectrum of biomarkers and imaging data in the *same* individuals. To overcome at least some of these shortcomings we generated genome-wide single-nucleotide polymorphism (SNP) genotyping data in the European Medical Information Framework Alzheimer’s Disease Multimodal Biomarker Discovery (EMIF-AD MBD) sample [11]. This powerful and unique dataset allows to combine genomic data (and “-omics” data from other domains) with preclinical biomarker levels to eventually improve our ability for an early detection and prevention of AD. While similar to the Alzheimer’s Disease Neuroimaging Initiative (ADNI) study [12] in various aspects, EMIF-AD MBD extends ADNI and scope in several important ways, e.g. in the breadth of the biomarker assessments as well as the availability of “-omics” data from various different domains in the same individuals (for more details see Bos et al., 2018 [11]).

In this report we focus exclusively on the description of the results from genome-wide association analyses using various Aβ and tau-relevant outcomes available in EMIF-AD MBD. Specifically, we performed GWAS and polygenic risk score (PRS) assessments for more than a dozen binary and quantitative phenotypes related to measures of cerebrospinal fluid (CSF) Aβ and tau proteins in addition to using simple diagnostic status (i.e. AD, MCI and control). Whenever available, we compare our findings using equivalent GWAS results from the ADNI dataset.

## Materials and Methods

### Sample and phenotype description

Overall, the EMIF-AD MBD dataset comprises 1221 elderly individuals (years of age: mean = 67.9, SD = 8.3) with different cognitive diagnoses at baseline (NC = normal cognition; MCI = mild cognitive impairment; AD = AD-type dementia). In addition, Aβ status, cognitive test results and at least two of the following were available at baseline for analyses in all EMIF-AD MBD individuals: plasma (n = 1189), DNA (n = 929), magnetic resonance imaging (MRI; n = 862), or CSF (n = 767 individuals). Furthermore, clinical follow-up data were available for 759 individuals. The demographic information of the 16 outcome phenotypes (9 binary and 7 quantitative) of the EMIF-AD MBD dataset utilized in this paper is summarized in Table 1. Depending on the availability of the clinical records, each phenotype has different effective sample sizes. We categorized the phenotypes analyzed in this study into three main categories, i.e. “diagnosis”, “amyloid protein assessment” and “tau protein assessment” (NB: the diagnostic criteria used here for AD are not incorporating on biomarker status so that some AD cases were classified as “amyloid negative”). Details related to sample ascertainment and phenotype / biomarker collection in EMIF-AD MBD have been described previously [11] and are summarized for the relevant traits of this study in Supplementary Table 1. Whenever available, we attempted to validate EMIF-AD MBD findings in the independent ADNI dataset using identical or comparable phenotypes (this was possible for the two diagnostic groups, as well as for 6 amyloid- and 2 tau-related traits; Table 1).

**Table 1.** Overview of binary and quantitative traits available for genome-wide association study (GWAS) and polygenic risk score (PRS) analyses in EMIF-AD MBD and ADNI datasets.

| Category | Variable type | Variable | Description | EMIF-AD (Discovery) |  |  | ADNI (Replication) |  |  |
| --- | --- | --- | --- | --- | --- | --- | --- | --- | --- |
|  |  |  |  | #sample_n | GWAS | PGS | Variable | #sample_n | GWAS |
| Clinical_diagnosis | binary | AD vs NC | Alzheimer disease (AD) vs normal cognition (NC) | 545 (212 AD vs 333 NC) | YES | YES | AD vs NC | 303 (46 AD vs 257 NC) | YES |
|  |  | MCI vs NC | Mild cognitive impairment (MCI) vs normal cognition (NC) | 659 (326 MCI vs 333 NC) | YES | YES | MCI vs NC | 705(448 MCI vs 257 NC) | YES |
| Amyloid_protein_assessment | binary | AMYLOIDstatus_ALL | Dichotomous amyloid classification variable across all diagnostic groups | 871 (455 abnormal vs 416 normal) | YES | YES | AMYLOIDstatus | 618 (361 abnormal vs 257 normal) | YES |
|  |  | AMYLOIDstatus_MCI | Dichotomous amyloid classification variable in MCI subjects | 326 (189 abnormal vs 137 normal) | YES | YES | AMYLOIDstatus_MCI | 371 (232 abnormal vs 139 normal) | YES |
|  |  | AMYLOIDstatus_NC | Dichotomous amyloid classification variable in NC subjects | 333 (77 abnormal vs 256 normal) | YES | YES | AMYLOIDstatus_NC | 202 (88 abnormal vs 114 normal) | YES |
|  |  | Central_CSF_ratiodich | Dichotomous variable based on ratio of central CSF amyloid-42/40 values | 677 (418 abnormal vs 259 normal) | YES | YES | NA | NA | NA |
|  |  | Local_AB42_Abnormal | Dichotomous variable of local CSF amyloid beta-42 values | 726 (392 abnormal vs 334 normal) | YES | YES | AB42_abnormal | 578 (340 abnormal vs 238 normal) | YES |
|  | quantitative | AB_Zscore | Z-score for amyloid pathology | 890 | YES | YES | AB_Zscore | 578 | YES |
|  |  | log_Central_CSF_AB42 | Log-transformed central CSF amyloid beta-42 values | 677 | YES | YES | ABETA.Lumi.bl | 578 | YES |
|  |  | Central_CSF_AB38 | Central CSF Amyloid beta-38 values | 675 | YES | YES | ABETA38.MSM.bl | 548 | YES |
|  |  | Central_CSF_AB40 | Central CSF Amyloid beta-40 values | 677 | YES | YES | ABETA40.MSM.bl | 548 | YES |
|  |  | log_Central_CSF_AB4240ratio | Ratio of log-transformed central CSF amyloid beta-42 vs. 40 values | 677 | YES | YES | NA | NA | NA |
| Tau_protein_assessment | binary | Local_TTAU_Abnormal | Dichotomous variable of local CSF total-tau values | 724 (378 abnormal vs 346 normal) | YES | YES | Not required | NA | NA |
|  |  | Local_PTAU_Abnormal | Dichotomous variable of local CSF phospho-tau values | 726 (354 abnormal vs 372 normal) | YES | YES | Not required | NA | NA |
|  | quantitative | Ttau_ASSAY_Zscore | Z-score for CSF total-tau values | 723 | YES | YES | log_TTAU_Zscore | 571 | YES |
|  |  | Ptau_ASSAY_Zscore | Z-score for CSF phospho-tau values | 726 | YES | YES | log_PTAU_Zscore | 576 | YES |

Replication data used in the preparation of this article were obtained from the ADNI database (adni.loni.usc.edu). The ADNI was launched in 2003 as a public-private partnership, led by Principal Investigator Michael W. Weiner, MD. The primary goal of ADNI has been to test whether serial MRI, positron emission tomography (PET), other biological markers, and clinical and neuropsychological assessment can be combined to measure the progression of MCI and early AD.

### DNA extraction

Our laboratory at University of Lübeck, Germany, had access to 953 DNA samples from EMIF-AD MBD participants [11] for genetic (this paper) and epigenetic (DNA methylation profiling, m.s. in preparation) experiments. All participants had provided written consent to these experiments and institutional review board (IRB) approvals for the utilization of the DNA samples in the context of EMIF-AD MBD were obtained by the sample collection sites. For 805 participants, DNA was extracted locally at the collection sites. For 148 whole blood samples DNA extraction was performed in our laboratory using the QIAamp® DNA Blood Mini Kit (QIAGEN GmbH, Hilden, Germany). Overall, this resulted in a total number of 953 DNA samples available for subsequent processing and analysis. Quality control (QC; by agarose gel electrophoresis, determination of A260/280 and A260/230 ratios, and PicoGreen quantification) resulted in 936 DNA samples of sufficient quality and quantity to attempt genome-wide SNP genotyping using the Infinium Global Screening Array (GSA) with Shared Custom Content (Illumina Inc.). GSA genotyping was performed at the Institute of Clinical and Medical Biology (UKSH, Campus-Kiel) on the iScan instrument (Illumina, Inc) following manufacturer’s recommendations. All 936 DNA samples passed post-experiment QC following the manufacturer’s instructions.

### Genotype imputation and quality control

Data processing was performed from raw intensity data (idat format) in GenomeStudio software (v2.0.4; Illumina, Inc.). We then used PLINK software (v1.9) [13] to perform pre-imputation QC and bcftools (v1.9) [14] to remove ambiguous SNPs, flipping and swapping alleles to align to human genome assembly GRCh37/hg19 before imputation. The QC’ed data (i.e. 931 samples and 498 589 SNPs) were then phased using SHAPEIT2 (v2.r837)[15] and imputed locally using Minimac3 [16] based on a precompiled Haplotype Reference Consortium (HRC) reference panel (EGAD00001002729 including 39 131 578 SNPs from ~11K individuals). Following post-imputation QC, we retained a total of 7 778 465 autosomal SNPs with minor allele frequency (MAF) ≥0.01 in 898 individuals of European ancestry for downstream association analysis. A full description of data processing and QC procedures is provided in the Supplementary Material.

### Classification of *APOE* genotypes

For all but 80 samples *APOE* genotype (i.e. for SNPs rs7412 [a.k.a. as “ɛ2-allele”] and rs429358 [a.k.a. “ɛ4-allele”]) was determined locally at the sample collection sites. To ensure that these prior genotypes correctly align to those resulting from genome-wide genotyping, local *APOE* genotypes were compared to those either inferred directly (i.e. rs7412) or indirectly (i.e. by imputation: rs429358) from GSA genotyping. These comparisons resulted in a total of 5 mismatches (~0.6%). In these 5 and the 80 samples without prior *APOE* genotype information, genotyping was determined manually in our laboratory using TaqMan assays (ThermoFisher Scientific, Foster City, CA) on a QuantStudio-12K-Flex system in 384-well format. TaqMan re-genotyping confirmed all 5 local genotype calls (which were henceforth used as genotypes in all subsequent analyses).

### Biochemical analyses of CSF biomarkers

CSF sampling and storage conditions have been described before [11]. CSF concentrations of Aβ38, Aβ40 and Aβ42 were measured using the V-PLEX Plus Aβ Peptide Panel 1 (6E10) Kit from Meso Scale Discovery (MSD, Rockville, MD). The measurements were performed at the Clinical Neurochemistry Laboratory in Gothenburg in one round of experiments, using one batch of kit reagents, by board-certified laboratory technicians, who were blinded to clinical data. For phosphorylated tau (Ptau) and total tau (Ttau), available data from the local cohorts were used. These were derived in clinical laboratory practice using INNOTEST ELISAs (Fujirebio, Ghent, Belgium), as previously described [17] In the absence of CSF for new analyses of CSF Aβ proteins, we used local INNOTEST ELISA-derived CSF Aβ42 data to allow classifying as many subjects as possible as either Aβ-positive or –negative (see [11] and below).

### GWAS and post-GWAS analyses

SNP-based association tests were performed using logistic regression models in mach2dat [18], [19] for binary traits and linear regression models in mach2qtl [18], [19] for quantitative traits. Association analyses utilized imputation-derived allele dosages as independent variables and were adjusted for sex, age at examination, and Principle component (PC) 1 to 5 (using PLINK –pca to compute eigenvalues for up to 20 PCs; the number of PCs was then determined visual inspection of the scree plot). Diagnostic groups (coded as AD = 3, MCI = 2, controls = 1) were included as additional covariates in all analyses except for diagnostic outcome. QQ and Manhattan plots were constructed in R version 3.3.3 (https://www.r-project.org) using the “qqman” package [20]. The genomic inflation factor was calculated in R using the “GenABEL” package [21]. Statistical significance for the SNP-based analyses was defined as α = 5E-08, a widely used threshold that accounts for the approximate number of independent variants (~1M) in European populations [22], [23]. Post-GWAS, we used FUMA (http://fuma.ctglab.nl/) [24] to perform functional mapping and annotation of the genome-wide association results. This included calculating gene-based association statistics using MAGMA [25] using predefined sets of genes as implemented in FUMA. Statistical significance for the gene-based analyses was defined as α=0.05/18720 = 2.671E-06 based on the number of genes (n=18720) utilized for these analyses, as suggested by FUMA [24].

### Polygenic risk score (PRS) analysis

PRS were calculated for each individual from the summary statistics of two partially overlapping AD case-control GWAS, i.e. the paper by Jansen et al. [7] including data from more than 380,000 individuals from the UK biobank, and a 2013 GWAS meta-analysis from the International Genomics of Alzheimer’s Project (IGAP) [8]. Note that the genome-wide screening data of the IGAP study (“stage 1”) was also included in the meta-analysis by Jansen et al.. After removal of ambiguous SNPs (A/T and C/G) and filtering SNPs by MAF >0.01 and imputation quality r2 >0.8, PLINK 1.9 software [13] was used for linkage disequilibrium (LD) pruning and scoring for a variety of P-value thresholds (5E-08, 5E-06, 1E-04, 0.01, 0.05, 0.10, 0.20, 0.30, 0.40, 0.50, 1.00). The resulting PRSs were used as independent variable in the regression models adjusting for sex, age, and PC1 to PC5 as covariates. For the phenotypes not representing the diagnostic outcome, we also included diagnosis as additional covariate. For linear models, variance explained (R^2^) was derived from comparing results from the full model (including PRS and covariates) vs the null model (linear model with covariates only). For logistic models, we calculated Nagelkerke’s r2 using the R package fmsb. A full description of these and all other statistical procedures is provided in the “Supplementary Methods”.

### Validation analyses in ADNI

Whenever possible, we used whole genome sequencing data from the ADNI cohort to assess replicability of the EMIF-AD MBD findings. The ADNI sample used here comprises 808 subjects with available whole genome sequencing data (for more details on the generation of these data, see http://adni.loni.usc.edu/study-design/). For the analyses performed here, we only used unrelated subjects of European origin (n=751). Variant-based filtering was performed based on minor allele count (MAC > 3), missingness rate (not more than 5 %) and Hardy-Weinberg equilibrium (p > 1E-05). We calculated principal components to account for population stratification based on an LD-pruned subset of common variants (MAF > 0.1). Association statistics were calculated using PLINK v2.0 using linear and logistic regression models (as appropriate), controlling for age, sex and four principal components as basic covariates in our models. For the phenotypes not representing the diagnostic outcome, we included diagnosis as an additional covariate. For more information on ADNI, please see Supplementary Material.

## Results

### GWAS on diagnostic outcomes

First, we performed GWAS for diagnostic outcomes, i.e. all cases diagnosed with AD (n=212) and MCI (n=326), against normal control subjects (n=333). Comparing AD vs. controls showed the expected strong signals in the *APOE* region on chromosome 19q reaching genome-wide significance in the gene-based tests (*APOC1* [P = 3.24E-07] and *APOE* [P = 3.39E-07]; Suppl. Figure 1b and Suppl. Table 2) and genome-wide suggestive association in the SNP-based analyses (best SNP rs429358: OR = 2.26, 95% CI = 1.66–3.07, P = 2.68E-07; Suppl. Figure 1a and Suppl. Table 2). Interestingly, and in contrast to most other previous AD GWAS (e.g. REFs for Lambert and /or Jansen), the best-associated SNP (rs429358) in this region is the variant defining the “ɛ4” allele in the commonly used “ɛ/2/3/4” haplotype (the “ɛ2” allele is defined by rs7412). *APOE* was also the top-associated region in the ADNI dataset (Suppl. Table 2), as previously described [26]. Interestingly, in analyses comparing MCI vs. controls, the *APOE* region did not emerge as strongly associated (i.e. P-value for rs429358 = 0.17; Suppl. Figure 2a/b, Suppl. Table 3). Instead, the best associated SNP was rs153308 (OR = 0.51, 95% CI = 0.39–0.67, P = 1.25E-06; Suppl. Figure 2a, Suppl. Table 3), located in an intergenic region on chromosome 5q13.3. However, neither this nor any of the other variants showing P-values <1E-05 showed any evidence of association in the ADNI dataset, thus they may – at least in part – not reflect genuine association signals.

### GWAS on dichotomous “amyloid classification”

In the remainder of our analyses we focus on AD-relevant CSF biomarker data as phenotypic outcomes of our GWAS analyses (see Table 1 for an overview of all traits analyzed). These analyses included a dichotomous “amyloid classification” (normal/abnormal) variable, representing a combination of CSFAβ42/40 ratio, local CSF-Aβ42, and the standardized uptake value ratio (SUVR) on an amyloid PET scan (see methods section on “amyloid classification” in Bos et al. [11] for more details). Using this amyloid variable (available for n=871 individuals, of which n=455 were classified as “abnormal” and n=416 as “normal”) across all diagnostic groups, we observed multiple genome-wide significant signals in the *APOE* region on chromosome 19 (Figure 1). Similar to the GWAS on diagnostic outcome, the strongest association was observed with the *APOE* “ɛ4” allele (i.e. SNP rs429358: OR = 5.66, 95% CI = 4.09- 7.84, P = 1.95E-25; Figure 1a, Suppl. Table 4), which was also very strong in equivalent analyses in the ADNI dataset (P = 4.15E-22). Despite the consistency of the *APOE* findings, none of the other suggestive signals outside the *APOE* region replicated in ADNI (Suppl. Table 4). As expected, gene-based tests using MAGMA highlighted *APOE* (and neighboring loci) as the most significant gene(s) associated with the “amyloid classification” variable (P = 1.13E-18, Figure 1b, Suppl. Table 4). In addition, a second locus emerged at genome-wide significance in the gene-based analyses, i.e. fermitin family homolog 2 (*FERMT2*) (P = 1.03E-06) located on chromosome 14q22.1. While this gene was originally reported to represent an AD risk locus [8], this notion was not replicated in the much larger GWAS by Jansen et al. [7]. Furthermore, gene-based tests in ADNI did not reveal evidence for association between markers in *FERMT2* and the “amyloid classification” phenotype (P = 0.23) suggesting that this gene is at best marginally involved in determining variance of this trait. Additional analyses using the “amyloid classification” variable limited to MCI and control individuals revealed a similar picture as in the full dataset (see Supplementary Tables 5 and 6, respectively), i.e. the strongest association signals were observed for *APOE* (all replicated in equivalent analyses in ADNI). In addition, we identified a few suggestive non-*APOE* signals on other chromosomes, albeit none of these showed evidence for independent replication in ADNI.

**Figure 1.**
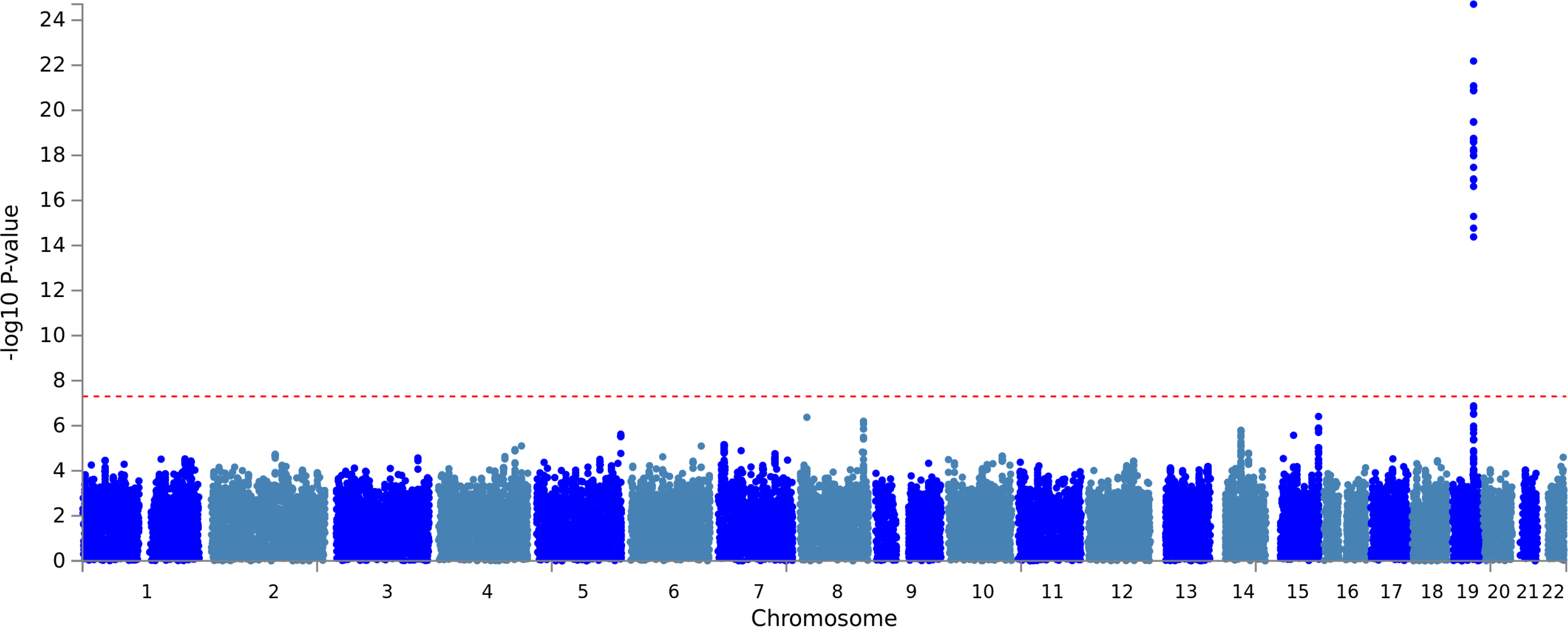
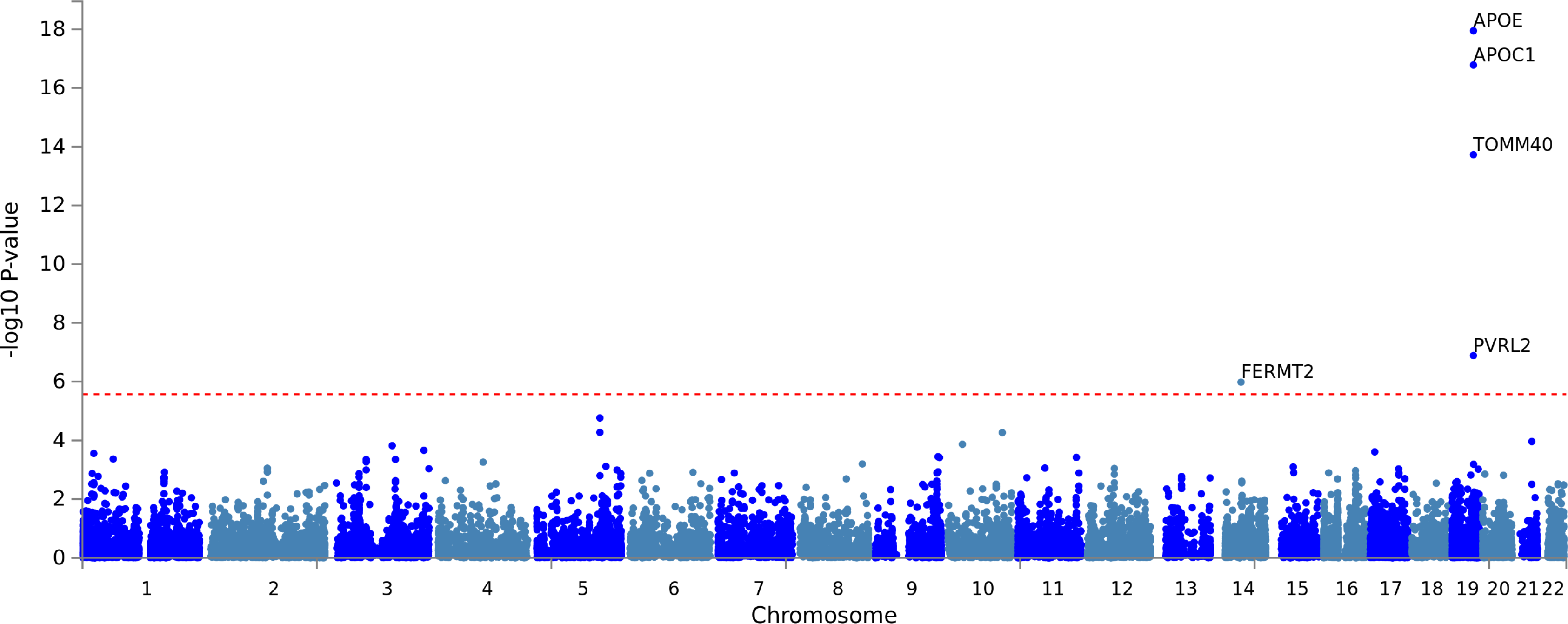
Manhattan plots of **1A)** SNP level and **1B)** gene-level genome-wide association results with the “amyloid classification” variable across all diagnostic groups (n = 871). All plots include gene assignments made with FUMA. Dotted red lines represent the threshold for genome-wide significance, i.e. α = 5.0E-08 for SNP-based (1A) and α = 2.671E-06 for gene-based (1B) analyses.

### GWAS on additional dichotomous and continuous CSF amyloid variables

Overall, there were a total of two binary (“Central_CSF_ratiodich” and “Local_AB42_Abnormal”) and five quantitative (“AB_Zscore”, “Central_CSF_AB38”, “Central_CSF_AB40”, “log_Central_CSF_AB42” and “log_Central_CSF_AB4240ratio”; see Table 1) CSF-Aβ phenotypes available that were used as outcome variables in the GWAS. With the exception of “CSF-Aβ38” and “CSF-Aβ40” all showed strong and highly significant association with markers in the *APOE* region but no better than suggestive (P < 1E-05) signals in the remaining genome (see Suppl. Figures, 3a/b-7a/b and Supplementary Table 7-11). None of the non-*APOE*, suggestive signals replicated in the ADNI dataset for phenotypes with available data. GWAS results on “CSF-Aβ38” (Figure 2, Suppl. Figure 8a/b and Supplementary Table 12) and “CSF-Aβ40” (Suppl. Figure 9a/b and Supplementary Table 13) showed highly similar GWAS results owing to the – well established [27] – high correlation between both markers which we also observed in this dataset (Pearson’s r^2^= 0.96, p-value < 2.2E-16 across all samples with available data). The most noteworthy finding emerging from these analyses was genome-wide significant association between CSF-Aβ40 and markers in the gene encoding zinc finger homeobox 3 (*ZFHX3*) in gene-based analyses (gene-based P-value = 7.48E-08; best SNP P-value = 7.29E-08; Suppl. Figure 9a/b and Supplementary Table 13; similar results were obtained with CSF-Aβ38). Despite the highly significant and consistent association between CSF-Aβ40 and CSF-Aβ38 levels and *ZFHX3*, this finding was not replicated in ADNI (gene-based P-value = 0.77; Supplementary Table 13).

**Figure 2.**
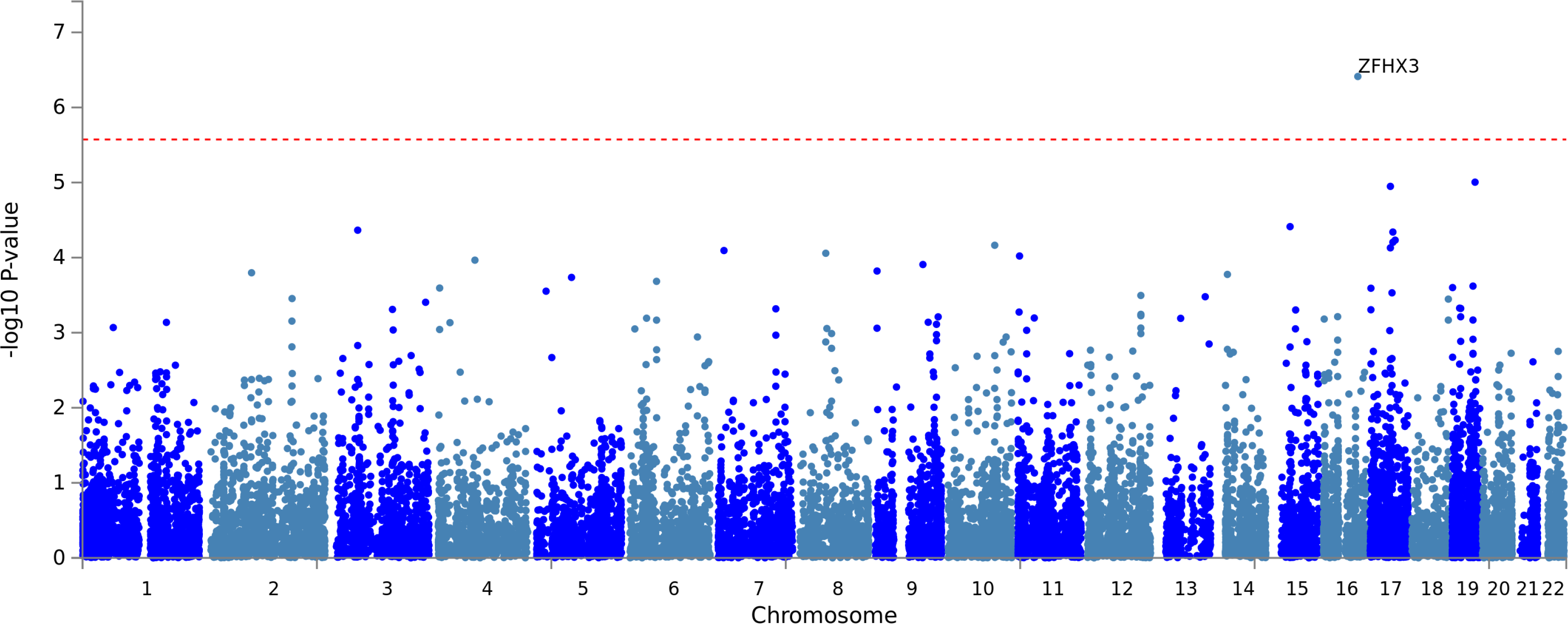
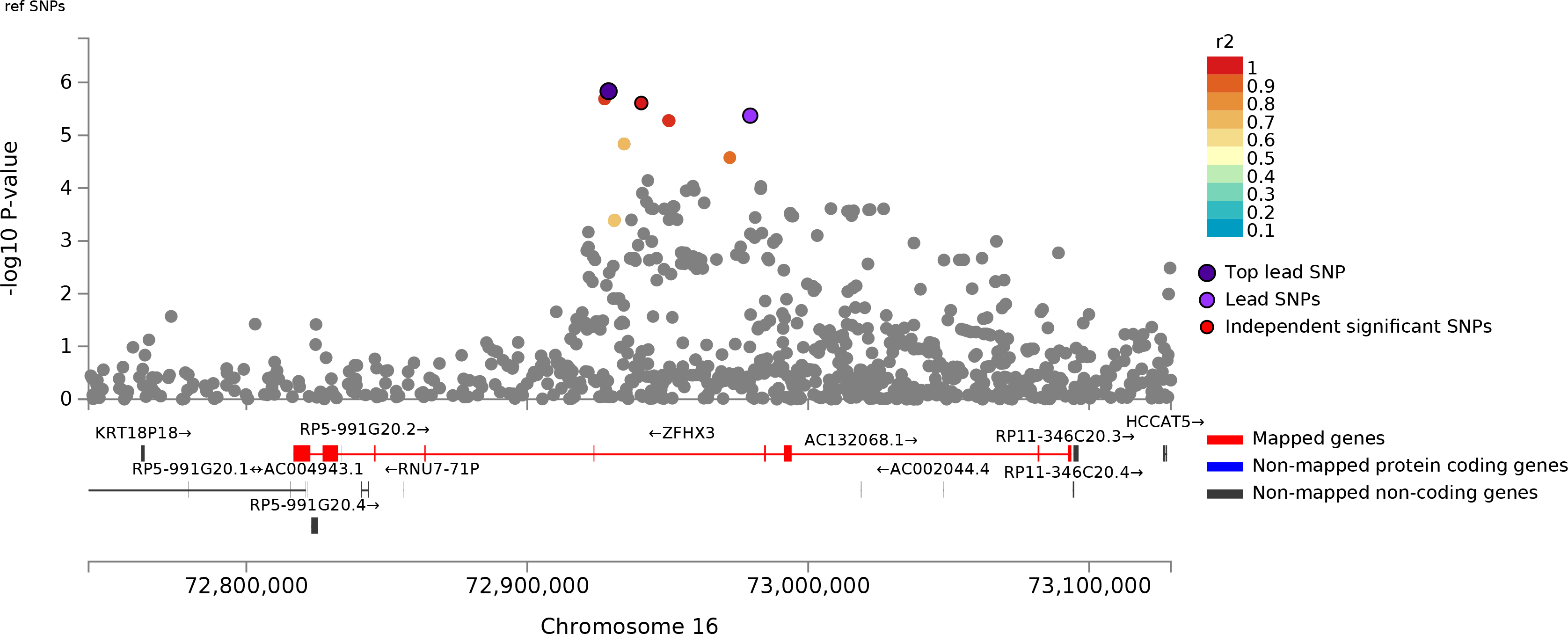
Manhattan plots of **2A)** gene-level genome-wide association results with CSF-Aβ38 levels across all diagnostic groups (n = 675), **2B)** regional association results zoomed into a 450kb region surrounding the *ZFHX3* gene on chromosome 16q22. All plots include gene assignments and linkage disequilibrium estimates made with FUMA. Dotted red line represents the threshold for genome-wide significance (α = 2.671E-06) for the gene-based analyses.

### GWAS on dichotomous and continuous CSF tau variables

Similar to the GWAS analyses for CSF Aβ measures, there were several dichotomous and continuous measures of CSF tau available in the EMIF-AD MBD dataset (see Table 1, Suppl. Figures 10a/b-13a/b). Using CSF-Ttau levels as z-scored continuous outcome we identified genome-wide significant association with markers in geminin coiled-coil domain containing (*GMNC*) on chromosome 3q28 in gene-based analyses (P = 1.61E-06; Suppl. Figure 11b, Suppl. Table 15). In the SNP-based analyses, markers in this gene showed genome-wide suggestive P-values ranging between 4.3E-06 and 9.5E-06 (Supplementary Table 15). These results showed marginal evidence for association in the ADNI cohort on the SNP level (for rs6444469 with P-value = 0.09), but not on the gene level (P-value = 0.59; Supplementary Table 15). Interestingly, association between markers in *GMNC* and CSF-Ttau levels were previously described [28]. A follow-up study to the original report provided evidence for sex-specific differences at this locus (rs1393060 proximal to *GMNC* and in strong LD with rs6444469 [r^2^=1]: P-value = 4.57E-10 in females compared to P-value = 0.03 in males) suggesting stronger effects in females [29]. In EMIF-AD MBD, we also observed a stronger association in females for *GMNC* at the gene and SNP level, respectively (gene: P-value = 4.09E-06 [in females] vs. P-value = 0.08 [in males]; SNP [for rs6444469]: P-value = 1.47E-04 [in females] vs. P-value = 8.91E-03 [in males]), hence, providing independent replication of the previous report. Similarly, the association results for rs6444469 and CSF-Ttau in ADNI were more pronounced in females (P-value = 0.013) than males (P-value = 0.93). The other available CSF tau measure was CSF-Ptau, which is known to strongly correlate with CSF-Ttau levels [30], a correlation which we also observe in our data (Pearson r^2^= 0.87, P-value < 2.2E-16). Owing to this phenotypic correlation, the GWAS results for both variables also look quite similar, as expected (Suppl. Figure 13b, Suppl. Table 17), and replicate the sex-specific difference.

### Polygenic risk score (PRS) analyses on all outcome traits

In addition to testing all above-mentioned traits for SNPs and genes in the context of genome-wide analyses, we also computed association statistics with aggregated variant data in the form of PRS using AD case-control results from Lambert et al. [8] and Jansen et al. [7]. The aim was to assess the degree at which established AD-associated markers also show association with the “AD-related” (endo-) phenotypes analyzed here. Although a considerable amount of sample overlap exists across the two AD risk GWAS (i.e. both use the “stage I” GWAS data from the IGAP sample in their discovery phase), both studies use different analysis paradigms and different replication cohorts. Given that the final effective sample size used in Jansen et al. is nearly 10-times larger than that in Lambert, we hypothesized that the results and, accordingly: PRS, derived from the larger study are more “precise” and will therefore show stronger association and explain more of the trait variance analyzed here. PRS-based results are summarized in Table 2, while full results can be found in Supplementary Table 18 (for PRS from Jansen et al. [7]) and 19 (for PRS from IGAP [8]). Overall, we observed significant PRS-based associations with many, but not all, traits analyzed in the EMIF-AD MBD dataset. For both PRS models, the best associated trait was a diagnosis of AD and all measures involving CSF-Aβ42 levels. In contrast, no noteworthy associations were observed with CSF-Aβ38 and CSF-Aβ40 levels nor with either of the two available CSF-tau measures (CSF-Ttau and CSF-Ptau). Comparing the “performance” of both PRS against each other revealed that – against our expectation - the IGAP-based results tended to show the stronger statistical support (i.e. smaller P-values) and explained slightly more of the phenotypic variance (i.e. showed higher r^2^) than the PRS derived from more recent and larger GWAS by Jansen et al. (Supp. Table 18 vs. 19). The only exception being the case-control analyses on AD status, where the Jansen PRS outperformed that from IGAP (i.e. r^2^ = 4.35%, P-value = 4.03E-06 vs. r^2^ = 2.98%, P-value = 1.07E-04, respectively). Interestingly, the association of AD-based PRS with risk for MCI was minor in both models (r^2^ = 0.83%, P-value = 0.02, and r^2^ = 0.65%, P-value = 0.039, for Jansen and IGAP, respectively).

**Table 2.** Summary of polygenic risk score (PRS) analyses using two P-value thresholds and two different GWAS datasets with and without markers in the *APOE* region. Red font = nominally significant association; bold red font = largest r^2^ (=most variance explained) for trait in question. A full listing of results from these PRS analyses can be found in Supplementary Table 18 (for Jansen et al GWAS) and Supplementary Table 19 (for IGAP GWAS).

| Phenotype | PRS constructed based on GWAS by Jansen et al, 2019 |  |  |  |  |  |  |  | PRS constructed based on GWAS by IGAP, 2013 |  |  |  |  |  |  |  |
| --- | --- | --- | --- | --- | --- | --- | --- | --- | --- | --- | --- | --- | --- | --- | --- | --- |
|  | Including *APOE* |  |  |  | Excluding *APOE* |  |  |  | Including *APOE* |  |  |  | Excluding *APOE* |  |  |  |
|  | S1(p<5E-8) |  | S5(<0.05) |  | S1(p<5E-8) |  | S5(<0.05) |  | S1(p<5E-8) |  | S5(<0.05) |  | S1(p<5E-8) |  | S5(<0.05) |  |
|  | R2* | Pvalue | R2 | Pvalue | R2* | Pvalue | R2 | Pvalue | R2* | Pvalue | R2 | Pvalue | R2* | Pvalue | R2 | Pvalue |
| AD | 3.09% | 8E-05 | 2.89% | 0.0002 | 1.46% | 0.0064 | 1.21% | 0.0131 | 2.98% | 0.0001 | 1.69% | 0.0034 | 0.78% | 0.0443 | 0.10% | 0.4745 |
| MCI | 0.12% | 0.3767 | 0.77% | 0.0249 | 0.06% | 0.5207 | 0.57% | 0.0532 | 0.51% | 0.0682 | 0.13% | 0.3537 | 0.49% | 0.0743 | 0.00% | 0.8764 |
| AMYLOIDstatus | 1.38% | 0.0004 | 1.01% | 0.0025 | 0.29% | 0.1056 | 0.05% | 0.4846 | 3.82% | 9E-09 | 2.95% | 3E-07 | 1.09% | 0.0018 | 0.34% | 0.0766 |
| Amyloid.MCI | 4.59% | 0.0008 | 5.82% | 0.0002 | 0.88% | 0.1377 | 0.84% | 0.1457 | 12.54% | 1E-07 | 7.45% | 2E-05 | 4.63% | 0.0008 | 0.82% | 0.1513 |
| Amyloid.NC | 0.83% | 0.1637 | 0.23% | 0.4657 | 0.20% | 0.4976 | 0.02% | 0.8144 | 2.45% | 0.0174 | 5.10% | 0.0008 | 0.24% | 0.4592 | 1.28% | 0.0866 |
| AB_Zscore | 1.20% | 0.0003 | 0.80% | 0.0031 | 0.22% | 0.1206 | 0.03% | 0.5801 | 2.61% | 7E-08 | 3.08% | 4E-09 | 0.39% | 0.0382 | 0.38% | 0.04 |
| Central_CSF_ratiodich | 3.86% | 2E-06 | 1.94% | 0.0007 | 1.29% | 0.0055 | 0.12% | 0.3945 | 5.87% | 9E-09 | 3.52% | 6E-06 | 1.58% | 0.0022 | 0.22% | 0.2439 |
| Local_AB42_Abnormal | 3.68% | 4E-06 | 1.13% | 0.0098 | 1.50% | 0.003 | 0.04% | 0.6249 | 3.40% | 1E-05 | 1.66% | 0.0018 | 0.94% | 0.0184 | 0.01% | 0.7873 |
| Central_CSF_AB38 | 0.21% | 0.2273 | 0.00% | 0.8587 | 0.21% | 0.2283 | 0.00% | 0.8638 | 0.00% | 0.8816 | 0.28% | 0.1614 | 0.03% | 0.6416 | 0.33% | 0.1279 |
| Central_CSF_AB40 | 0.24% | 0.1805 | 0.01% | 0.7974 | 0.18% | 0.2553 | 0.03% | 0.6317 | 0.05% | 0.5418 | 0.44% | 0.0718 | 0.07% | 0.4874 | 0.36% | 0.106 |
| log_Central_CSF_AB4240ratio | 2.06% | 4E-05 | 1.68% | 0.0002 | 0.41% | 0.0694 | 0.11% | 0.3518 | 3.74% | 3E-08 | 2.55% | 5E-06 | 0.56% | 0.0323 | 0.05% | 0.5324 |
| log_Central_CSF_AB42 | 1.52% | 0.0005 | 0.52% | 0.0445 | 0.41% | 0.075 | 0.00% | 0.9801 | 2.31% | 2E-05 | 2.17% | 4E-05 | 0.53% | 0.0425 | 0.25% | 0.1605 |
| Local_PTAU_Abnormal | 0.31% | 0.1602 | 0.03% | 0.662 | 0.11% | 0.3957 | 0.00% | 0.9458 | 0.69% | 0.0347 | 0.38% | 0.1192 | 0.19% | 0.2697 | 0.09% | 0.4424 |
| Local_TTAU_Abnormal | 0.93% | 0.0107 | 0.01% | 0.7505 | 0.66% | 0.0305 | 0.13% | 0.3353 | 0.82% | 0.0166 | 0.27% | 0.1644 | 0.17% | 0.267 | 0.04% | 0.5771 |
| Ptau_ASSAY_Zscore | 0.34% | 0.0909 | 0.00% | 0.9833 | 0.19% | 0.2121 | 0.01% | 0.7372 | 0.37% | 0.0806 | 0.08% | 0.4258 | 0.01% | 0.7755 | 0.01% | 0.8237 |
| Ttau_ASSAY_Zscore | 0.51% | 0.032 | 0.03% | 0.6082 | 0.32% | 0.0909 | 0.00% | 0.9538 | 0.09% | 0.3566 | 0.11% | 0.3269 | 0.03% | 0.5811 | 0.00% | 0.8676 |

To investigate the contribution of markers in the *APOE* region, we repeated all analyses excluding variants within 1 MB of *APOE* (chr19:45409039-45412650; bottom part of Supp. Tables 18 and 19). As expected, removal of *APOE* region markers from the PRS decreased the variance explained for most of the traits analyzed here, albeit to varying degrees. Most affected were the analyses on AD (strongest reduction in r^2^ in analyses excluding *APOE* effects vs. the full model including *APOE* = 73.8% in IGAP) and essentially all CSF-Aβ42 related measures (strongest reduction in r^2^ = 88.9% for trait “AB_Zscore” in IGAP) for both PRS models. Less affected by the removal of *APOE* were the analyses of CSF-tau species (strongest reduction in r^2^ = 29.0% for CSF-Ttau in non-*APOE* vs. *APOE* models).

## Discussion

This is the first GWAS utilizing part of the wide array of AD-relevant phenotypes and biomarkers available in the EMIF-AD MBD dataset. The phenotypes analyzed here related either to clinical diagnosis (i.e. AD or MCI) or to levels of CSF biomarkers revolving around various biochemical species of amyloid or tau proteins. While GWAS results have already been reported for some of the biomarkers analyzed here (e.g. in ADNI), ours are the first to combine genomic and biomarker data in the newly established EMIF-AD MBD dataset. The main findings of our study can be summarized in the following five points: 1) the most prominent genetic signals in analyses of either diagnostic outcome or phenotypes related to CSF-Aβ42 were observed with markers in or near *APOE*, which is in good agreement with equivalent analyses in ADNI and other datasets; 2) our analyses identified one novel association in analyses of CSF-Aβ38 and CSF-Aβ40 levels and DNA sequence variants in *ZFHX3* (a.k.a. *ATBF1* [AT-motif binding factor 1]), although these signals were not replicated in the ADNI dataset; 3) using CSF-tau species (i.e. Ttau and Ptau), we confirmed the previously described association with SNPs in *GMNC*, including the recently reported effect modification by sex at this locus; 4) PRS analyses revealed that AD risk SNPs are mostly associated with phenotypes related to CSF-Aβ42 but not CSF-Aβ38 or CSF-Aβ40 or CSF-tau; 5) exclusion of *APOE* from the PRS analyses suggest that non-*APOE* AD GWAS SNPs explain at most 2.5% of the phenotypic variance underlying a diagnosis of AD in this dataset. Collectively, these results implicate that the genetic architecture underlying many traits relevant for AD research in EMIF-AD MBD compare well to other datasets of European descent paving the way for future genomic discoveries with additional phenotypes available in this unique cohort [11].

Despite these promising first results, our study is potentially confined by a number of possible limitations: First and foremost, despite the breadth of available phenotype data, the overall sample size of the EMIF-AD MBD dataset is comparatively small and, as a result, may not be adequately powered to detect genetic variants exerting smaller effects. To a degree, this limitation is alleviated by the fact that many available outcome phenotypes are of a quantitative nature, which are more powerful than analyses of dichotomous traits (e.g. disease risk). Second, many of the phenotypes available in EMIF-AD MBD are not currently ascertained in other, independent datasets (such as ADNI), making independent replication of any novel findings difficult. This situation can be expected to improve somewhat once the phenotypic breadth in ADNI and other cohorts is extended. Still, until independent replication is available, novel GWAS findings from EMIF-AD MBD must be interpreted with caution. This includes the putative association between CSF-Aβ38 and CSF-Aβ40 and markers in *ZFHX3*, which were highly significant and consistent in EMIF-AD MBD but not replicated in ADNI. Until more independent data on these outcome traits are available, the *ZFHX3* association should be considered “provisional”. In this context it is comforting, however, that many well-established genetics findings (such as the association between *APOE* and measures of CSF-Aβ or *GMNC* and CSF-tau) were reproduced in EMIF-AD MBD. Third, while the genome-wide SNP genotype data was generated in one run of consecutive experiments in one laboratory, the same is not true for the phenotype measurements, which were performed locally in each of the 11 participating sites. In some instances, the compiled phenotype data are not based on the same biochemical assays across sites for some variables, e.g. measurements of tau protein. While the EMIF-AD MBD phenotype team went to great lengths to alleviate this potential problem by normalizing variables for each center (see Bos et al. [11] for more details), the possibility of artefactual findings owing to phenotypic heterogeneity remains. Finally, as described in the overall cohort description manuscript, the EMIF-AD MBD dataset is not designed to be “representative” of the general population but was assembled with the aim to achieve approximately equal proportions of amyloid+ vs. amyloid- individuals in all three diagnostic subgroups. While this ascertainment strategy does not invalidate our GWAS results per se, they may not be generalizable to the population as a whole. However, this limitation may affect any study with clinically ascertained participants and, thus, applies to most previously published GWAS in the field, including those performed in ADNI.

In conclusion, our first-wave of GWAS analyses in the EMIF-AD MBD dataset provides a first important step in a series of additional genome-wide and epigenome-wide (using DNA methylation profiles) association analyses in this valuable and unique cohort.

## Author contributions

SH performed all the analyses and interpretation on data and wrote the manuscript. DP and RT contributed to replication in ADNI. VD was responsible for EMIF-AD MBD DNA sample preparation and extraction. FK was responsible for EMIF-AD MBD genotypic data QC and imputation. IB, SV, BT, and PJ coordinated the collection and harmonization of phenotypes and biosamples in EMIF-AD MBD and helped identifying equivalent phenotypes from the ADNI catalog. AF and MW supervised the genotyping experiments. UA, KB and HZ performed CSF biomarker measurements and took part in cut-point determinations. KS and CVB contributed to genetic characterization of samples and design of the genomics studies in EMIF-AD MBD, and critically revised the manuscript drafts. RV, SG, JS and IC contributed to sample and data collection. EN and IS were responsible for data collection of the Antwerp cohort. AW and PK contributed to data and sample collection / handling of the Gothenburg MCI Study, Gothenburg, Sweden. JS, PJV and SL are leads for the EMIF-AD MBD; designed and managed the platform. LB designed and supervised the genomics portion of the EMIF-AD MBD project and co-wrote all drafts of the manuscript. All authors critically revised all manuscripts drafts, read and approved the final manuscript.

### Alzheimer’s Disease Neuroimaging Initiative

Data used in preparation of this article were obtained from the Alzheimer’s Disease Neuroimaging Initiative (ADNI) database (http://adni.loni.usc.edu). As such, the investigators within the ADNI contributed to the design and implementation of ADNI and/or provided data but did not participate in analysis or writing of this report. A complete listing of ADNI investigators can be found at: http://adni.loni.usc.edu/wp-content/uploads/how_to_apply/ADNI_Acknowledgement_List.pdf.

## Acknowledgements

The present study was conducted as part of the EMIF-AD MBD project, which has received support from the Innovative Medicines Initiative Joint Undertaking under EMIF grant agreement n° 115372, the resources of which are composed of financial contribution from the European Union’s Seventh Framework Program (FP7/2007–2013) and EFPIA companies’ in kind contribution. Parts of this study were made possible through support from the German Research Foundation (DFG grant FOR2488: Main support by subproject “INF-GDAC” BE2287/7-1 to LB). RV acknowledges the support by the Stichting Alzheimer Onderzoek (#13007, #11020, #2017-032) and the Flemish Government (VIND IWT 135043). HZ is a Wallenberg Academy Fellow supported by grants from the Swedish Research Council (#2018-02532), the European Research Council (#681712) and Swedish State Support for Clinical Research (#ALFGBG-720931). SJBV received funding from the Innovative Medicines Initiative 2 Joint Undertaking under ROADMAP grant agreement No. 116020 and from ZonMw during the conduct of this study. No conflict of interest exists. Research at VIB-UAntwerp was in part supported by the University of Antwerp Research Fund. The research was supported by ALF clinical grants from Region Västra Götaland to AW and to PK. The authors acknowledge the assistance of Ellen De Roeck, Naomi De Roeck and Hanne Struyfs (UAntwerp) with data collection. The Lausanne study was funded by a grant from the Swiss National Research Foundation (SNF 320030_141179) to JP. We thank Mrs. Tanja Wesse and Mrs. Sanaz Sedghpour Sabet at the Institute of Clinical Molecular Biology, Christian-Albrechts-University of Kiel, Kiel, Germany for technical assistance with the GSA genotyping. The LIGA team acknowledges computational support from the OMICS compute cluster at the University of Lübeck. The computations in the ADNI cohort were run on the Odyssey cluster supported by the FAS Division of Science, Research Computing Group at Harvard University.

For ADNI: Data was used for this project of which collection and sharing was funded by the Alzheimer’s Disease Neuroimaging Initiative (ADNI) (National Institutes of Health Grant U01 AG024904) and DOD ADNI (Department of Defense award number W81XWH-12-2-0012). ADNI is funded by the National Institute on Aging, the National Institute of Biomedical Imaging and Bioengineering, and through generous contributions from the following: AbbVie, Alzheimer’s Association; Alzheimer’s Drug Discovery Foundation; Araclon Biotech; BioClinica, Inc.; Biogen; Bristol-Myers Squibb Company; CereSpir, Inc.; Cogstate; Eisai Inc.; Elan Pharmaceuticals, Inc.; Eli Lilly and Company; EuroImmun; F. Hoffmann-La Roche Ltd and its affiliated company Genentech, Inc.; Fujirebio; GE Healthcare; IXICO Ltd.; Janssen Alzheimer Immunotherapy Research & Development, LLC.; Johnson & Johnson Pharmaceutical Research & Development LLC.; Lumosity; Lundbeck; Merck & Co., Inc.; Meso Scale Diagnostics, LLC.; NeuroRx Research; Neurotrack Technologies; Novartis Pharmaceuticals Corporation; Pfizer Inc.; Piramal Imaging; Servier; Takeda Pharmaceutical Company; and Transition Therapeutics. The Canadian Institutes of Health Research is providing funds to support ADNI clinical sites in Canada. Private sector contributions are facilitated by the Foundation for the National Institutes of Health (http://www.fnih.org). The grantee organization is the Northern California Institute for Research and Education, and the study is coordinated by the Alzheimer’s Therapeutic Research Institute at the University of Southern California. ADNI data are disseminated by the Laboratory for Neuro Imaging at the University of Southern California.

## Additional files

The Supplement to this manuscript contains Supplementary Methods, Supplementary Figures 1-13 and Supplementary Tables S1-19.

