## Supplementary Material for "Genome-wide association study of Alzheimer’s disease CSF biomarkers in the EMIF-AD Multimodal Biomarker Discovery dataset"

ADNI started in 2003 as a public-private collaboration under the supervision of Principle Investigator Michael W. Weiner, MD. The primary goal of ADNI is to study whether serial magnetic resonance imaging (MRI), positron emission tomography (PET), other biological markers, and clinical and neuropsychological measures can be combined to measure the progression of mild cognitive impairment (MCI) and early Alzheimer’s disease (AD). Please see www.adni-info.org for the latest information.

### Genotype data handling, QC and imputation procedures

Raw data processing, i.e. clustering and genotype calling from raw intensity data (idat format) was performed in GenomeStudio software (Illumina, Inc.) using the genotyping module (version 2.0.2). Samples with call rate <0.95 and p50GC <0.7 were excluded at this stage. We then used PLINK software (**v1.9;** [1]**)** to perform additional QC filtering, i.e. sex checks (--check-sex 0.25 0.75), strand check (--flip), missing genotype rate (--geno 0.02; --mind 0.05), Hardy-Weinberg equilibrium (HWE) tests (--hwe 0.000005), and minor allele frequency (MAF) filtering (--maf 0.01). For determining pairwise allele sharing (to identify cryptic relatedness), we used an LD-pruned set of markers (--indep-pairwise 1500 150 0.2). Pairwise allele-sharing IBD/IBS was determined using (--Z-genome --min 0.1). Overall, this procedure led to 498,589 QC-filtered SNPs in 931 samples suitable for imputation. The LD pruned dataset was also used for principal component analysis (PCA; using PLINK command ‘--pca') along with the reference dataset of the 1000 Genome Project Consortium Phase 3 (The 1000 Genomes Project Consortium, 2015) to assign ethnic descent groups using the five 1000G super-populations by k-nearest neighbor (k-NN; k=9) classification (using R package ‘class’ in R 2.3.2; [2]). This resulted in assigning a “European descent” to 898 out of all 931 samples; only these n=898 samples were used in the subsequent statistical analyses.

Before imputation, the QC’ed genotype data were then subjected to bcftools (v1.9) [3] for removing ambiguous SNPs, and flipping and swapping alleles to align to GRCh37/hg19. This was followed by haplotype phasing using SHAPEIT2 [4] and imputation of unobserved genotypes using Minimac3 [5] using a precompiled Haplotype Reference Consortium (HRC) reference panel (EGAD00001002729 including 39,131,578 SNPs from ~11K individuals). Post-imputation we only retained autosomal SNPs with minimac3 Rsq ≥0.3, MAF ≥1%, and HWE P-values ≥ 5E-6 (using best-guess genotypes in control subjects), leaving a total of 7,778,465 SNPs for statistical analyses.

**Polygenic risk score (PRS) analysis**

To aggregate data on multiple variants per individual we computed polygenic risk scores (PRS) for each individual which were then used as independent variable in the statistical analyses. Allele status and effect-size estimates were taken from the summary statistics of the largest AD GWAS published to date [6] and compared to the 2013 GWAS results from IGAP [7]. First, we removed ambiguous SNPs (A/T and C/G) from the list of considered variants. Furthermore, we only used SNPs with MAF >0.01 and imputation quality r2 >0.8 in the EMIF-AD dataset. Next, LD pruning was performed on the CEU portion of the HRC reference panel used for imputation. To this end, we used PLINK 1.9 software [1] for two consecutive rounds of marker pruning (1$^{st}$ round: --clump-p1 1 --clump-p2 1 --clump-r2 .5 --clump-kb 250; 2$^{nd}$ round: --clump-p1 1 --clump-p2 1 --clump-r2 .2 --clump-kb 5000). Actual PRS were computed using the --score command in PLINK for a variety of P-value thresholds in the primary GWAS data (0.00000005, 0.000005, 0.0001, 0.01, 0.05, 0.10, 0.20, 0.30, 0.40, 0.50, 1.00). The resulting PRS were used as independent variable in the linear or logistic regression models adjusting for sex, age, and PC1 to PC5 as covariates. For the phenotypes not representing the diagnostic outcome, we also included diagnosis as additional covariate. For linear models, variance explained (r2) was derived from comparing results from the full model (including outcome phenotype and covariates) vs the null model (linear model with covariates only). For logistic models, we calculated Nagelkerke's $r^2$ using the R package fmsb.

**Supplementary Tables**

Supplementary Tables 1-19 can be found in the MS-Excel file "Supplementary_Tables_20190923.biorxiv.xls".

**Supplementary Figures**

**All supplementary figures are arranged in the same format ("X" stands for figure number). See Table 1 in the main text for a full description of variable names.**

Fig XA: Manhattan (top) and quantile-to-quantile (bottom) plots of **SNP-based** genome-wide association results. Red line represents the threshold for genome-wide significance ($\alpha$ = 5.0E-08).

Fig XB: Manhattan (top) and quantile-to-quantile (bottom) plots of **gene-based** genome-wide association results. Red line represents the threshold for genome-wide significance ($\alpha$ = 2.671E-06). Gene-based plots drawn in FUMA.

**Figure S1a.** AD vs NC

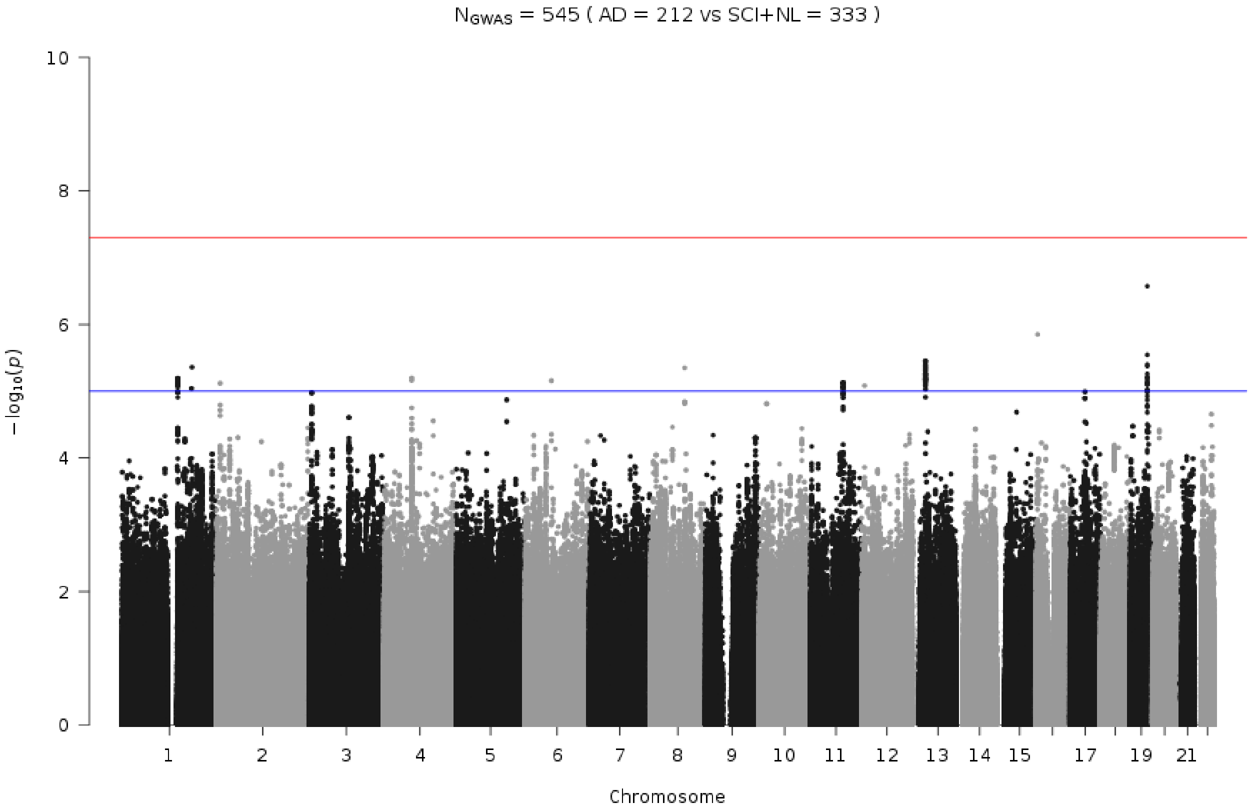


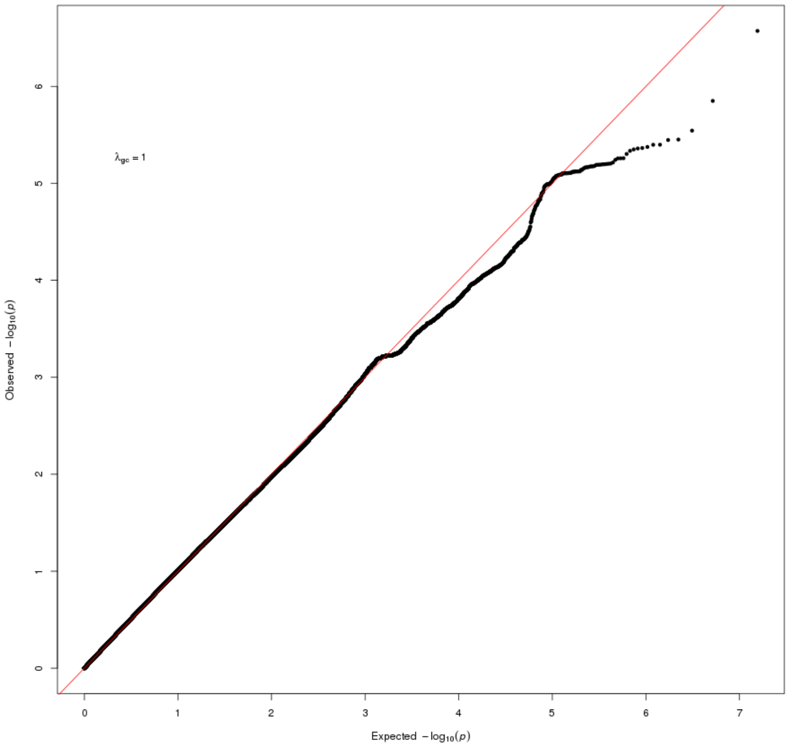

**Figure S1b.** AD vs NC

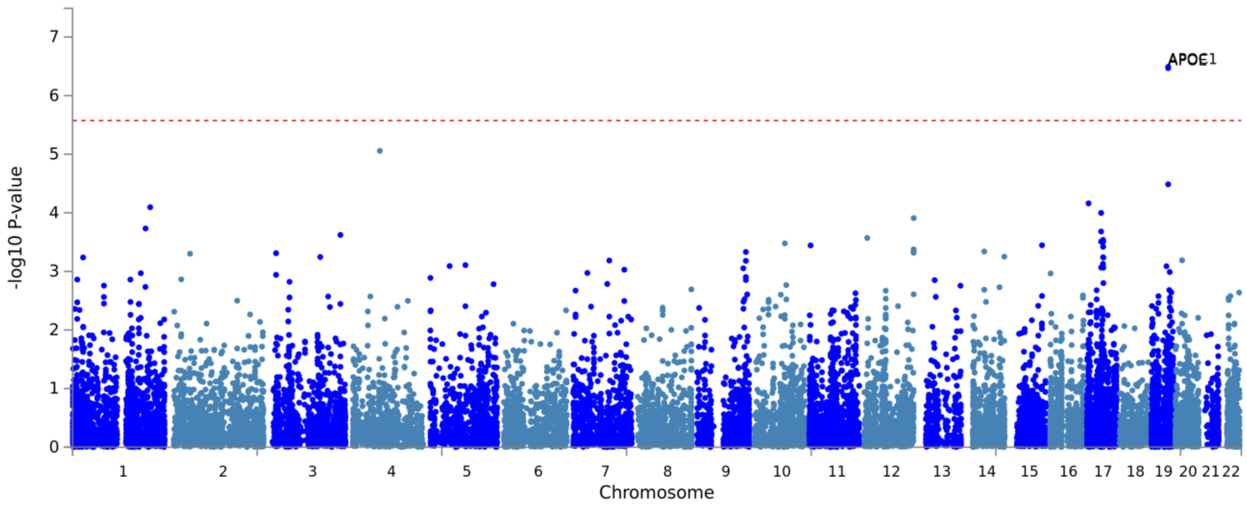


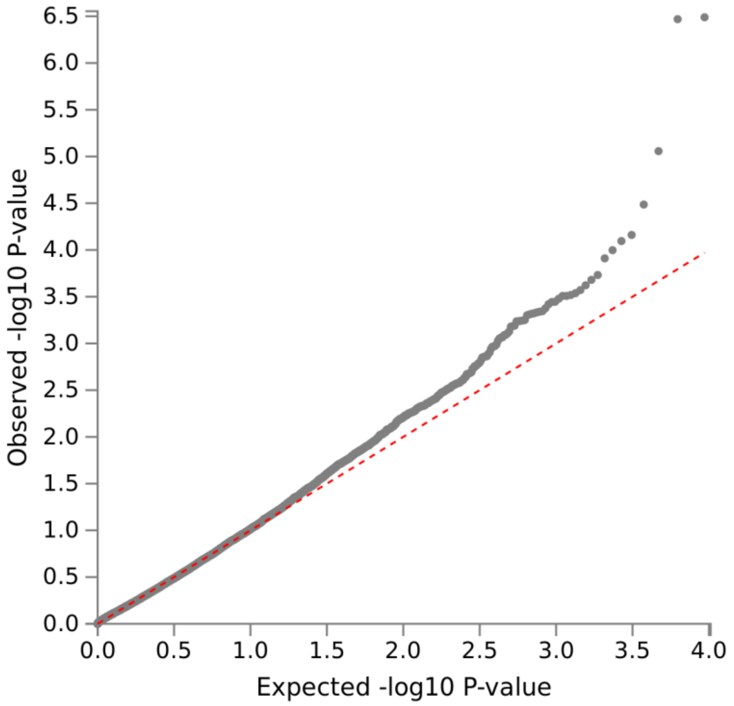

**Figure S2a.** MCI vs NC

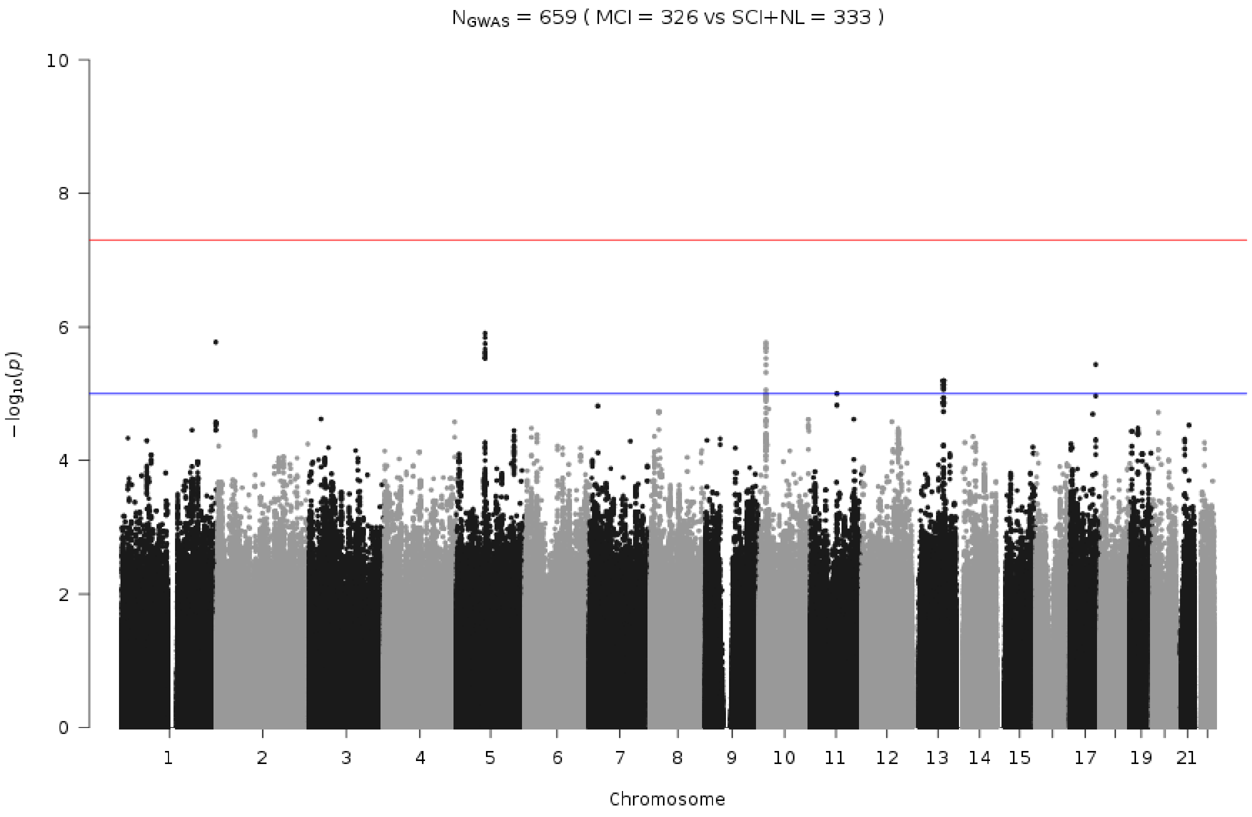


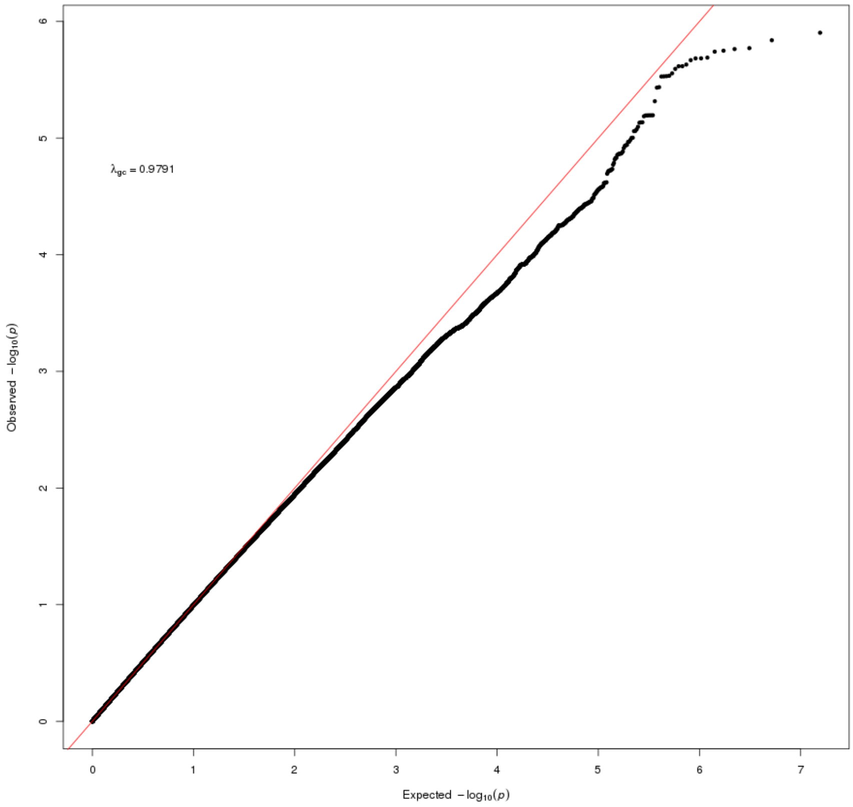

**Figure S2b.** MCI vs NC

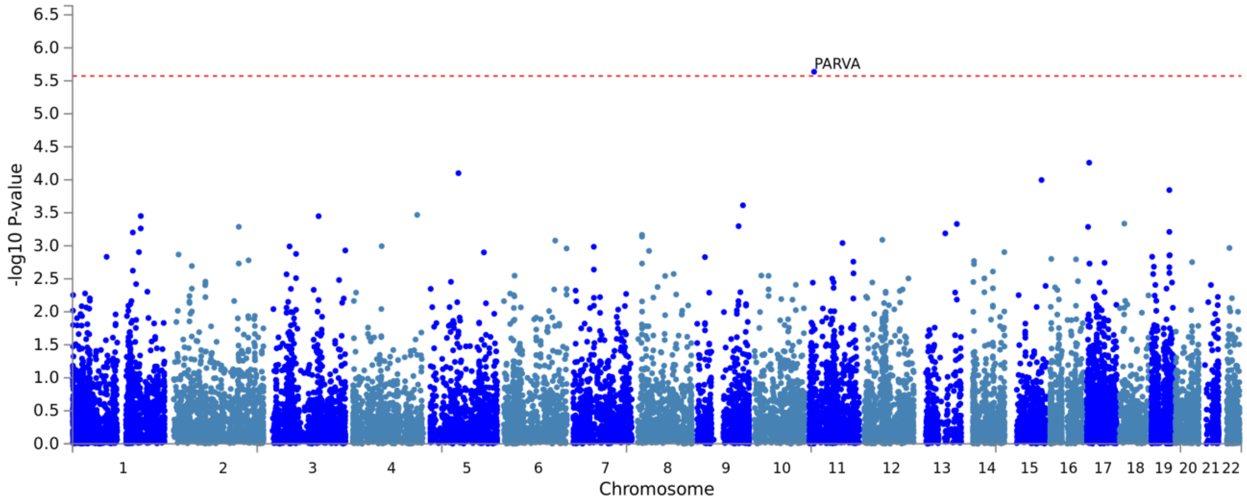


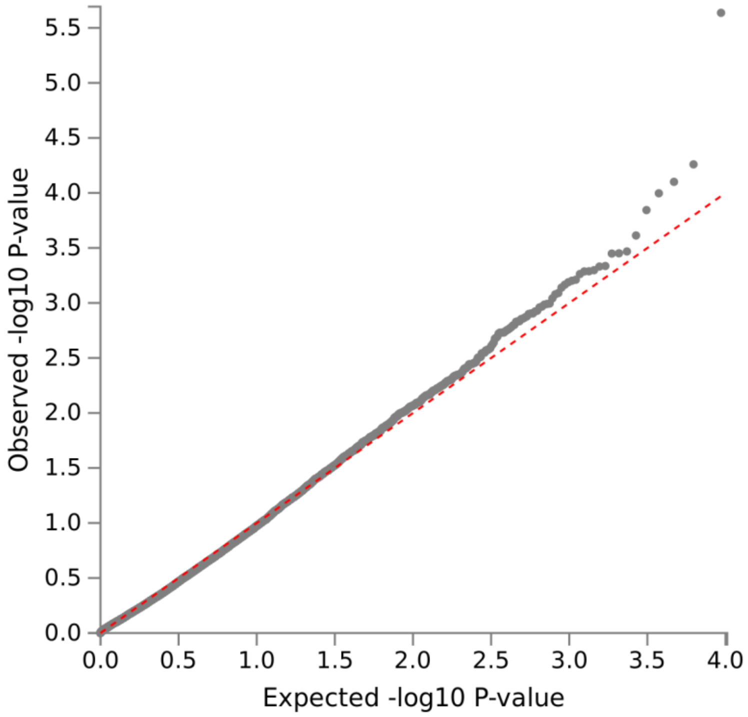

**Figure S3a.** Central_CSF_ratiodich

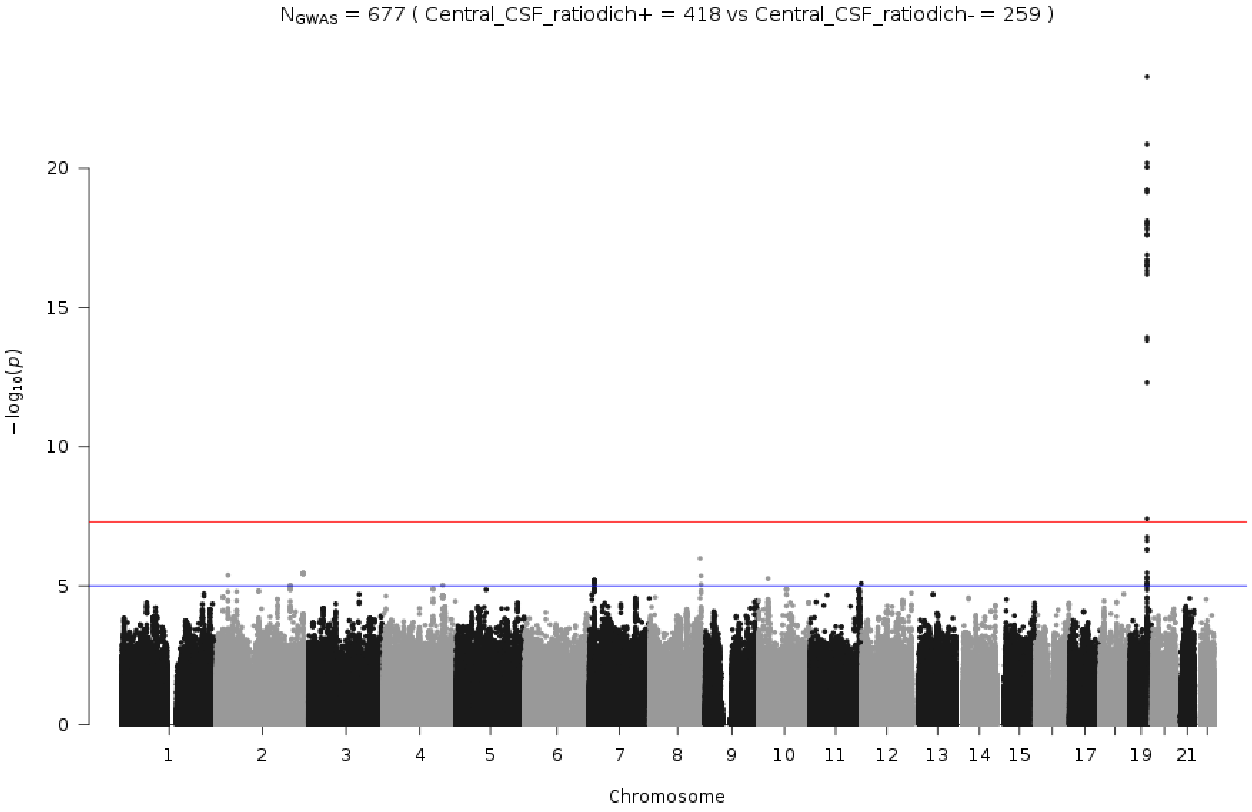


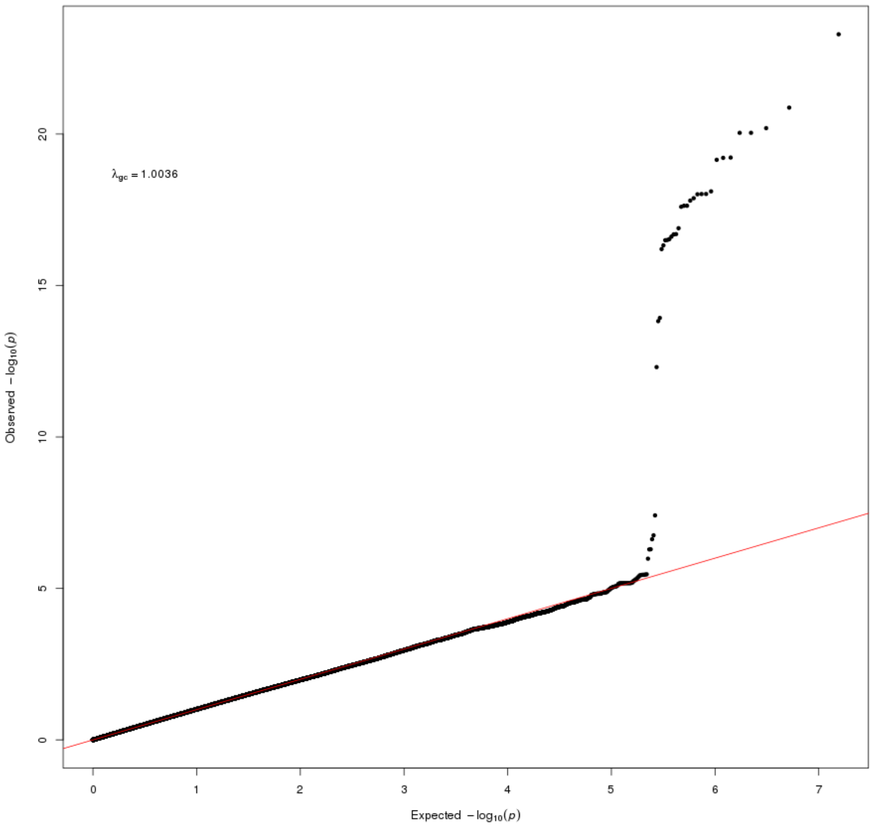

**Figure S3b.** Central_CSF_ratiodich

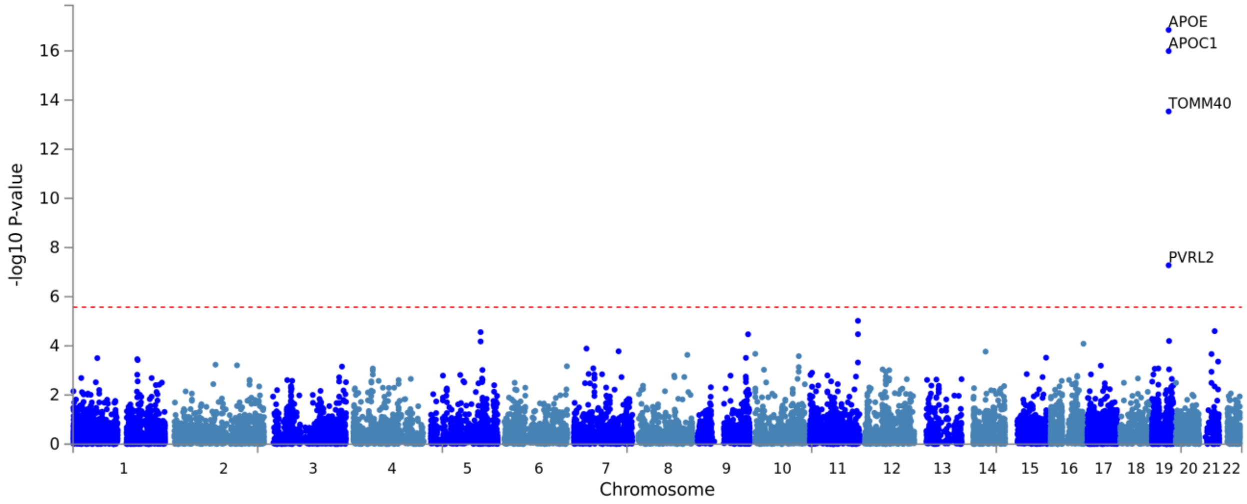


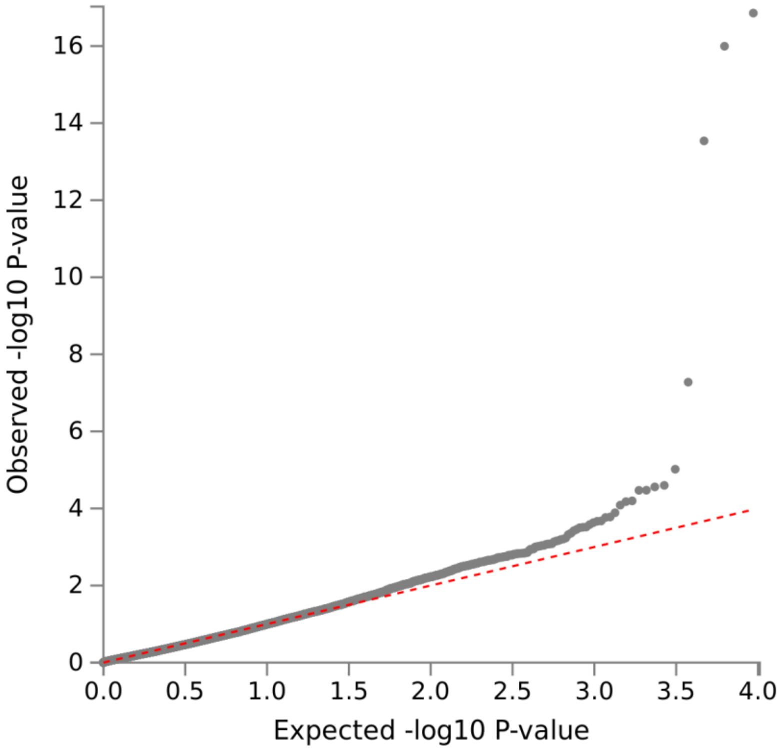

**Figure S4a.** Local_AB42_Abnormal

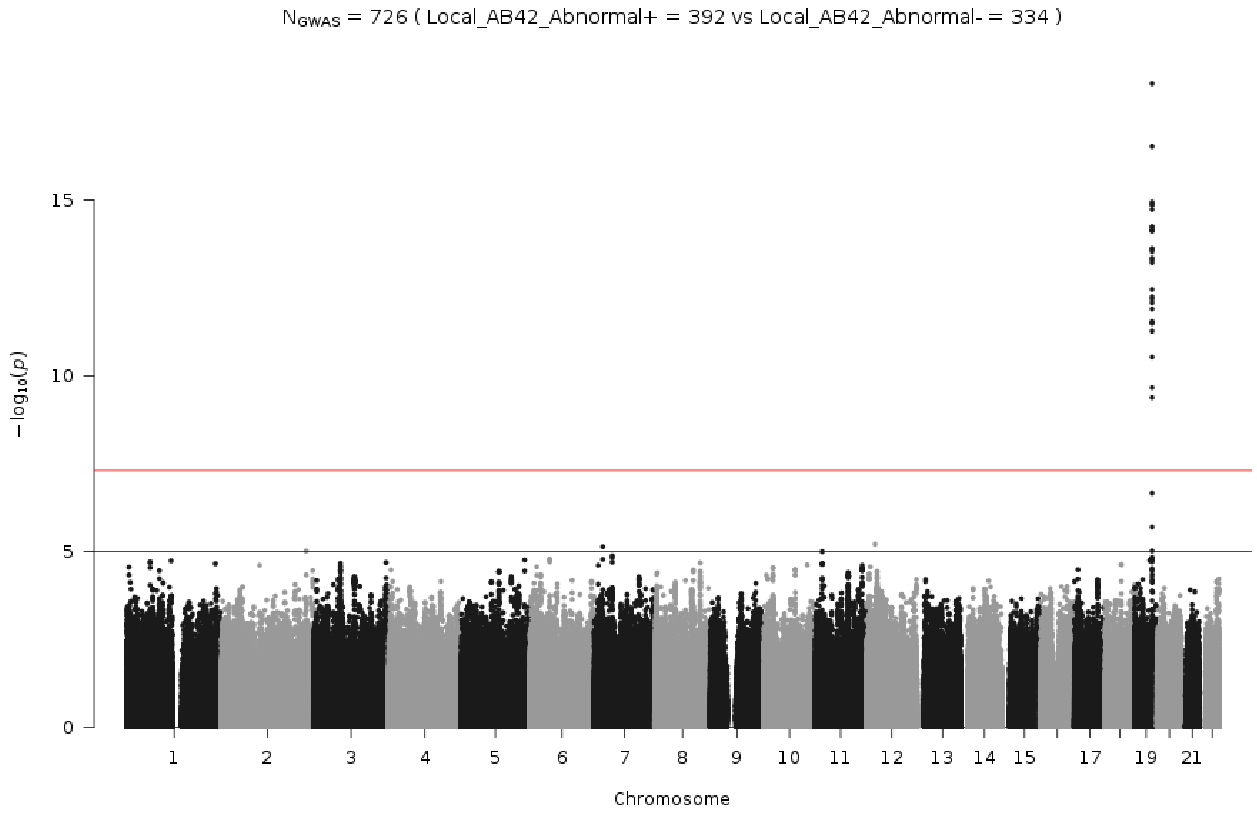


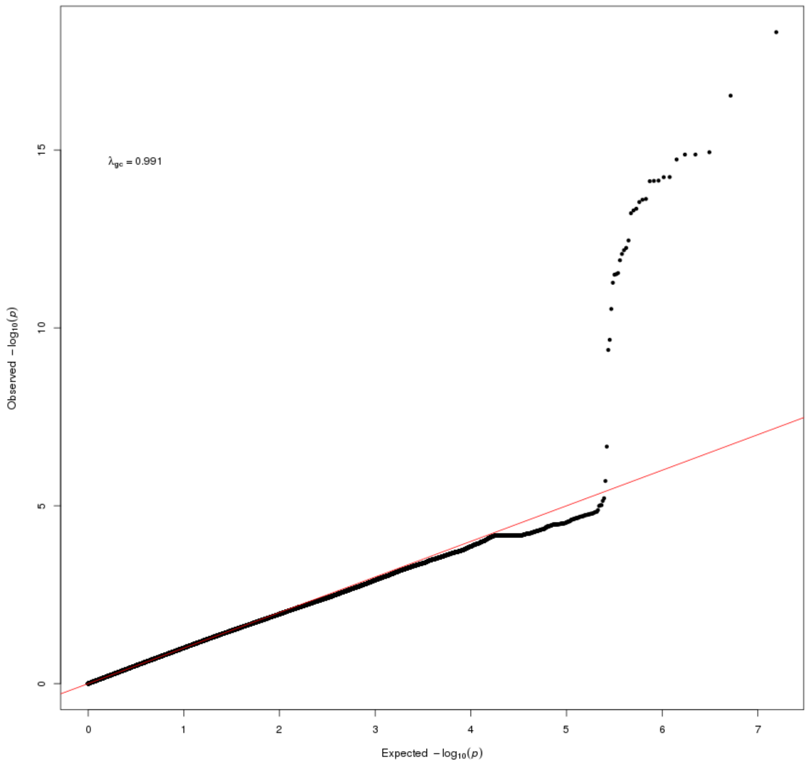

**Figure S4b.** Local_AB42_Abnormal

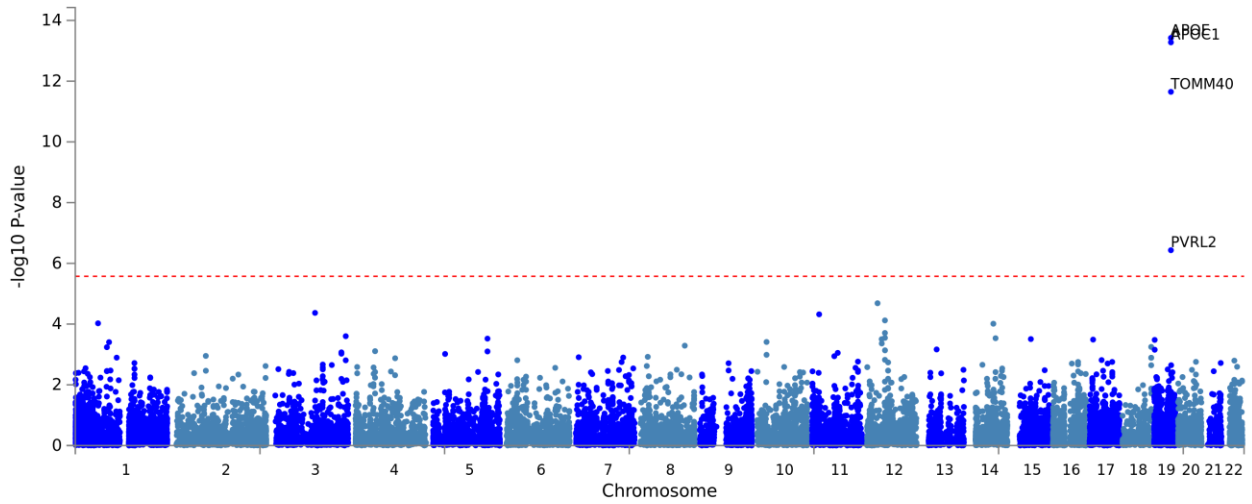


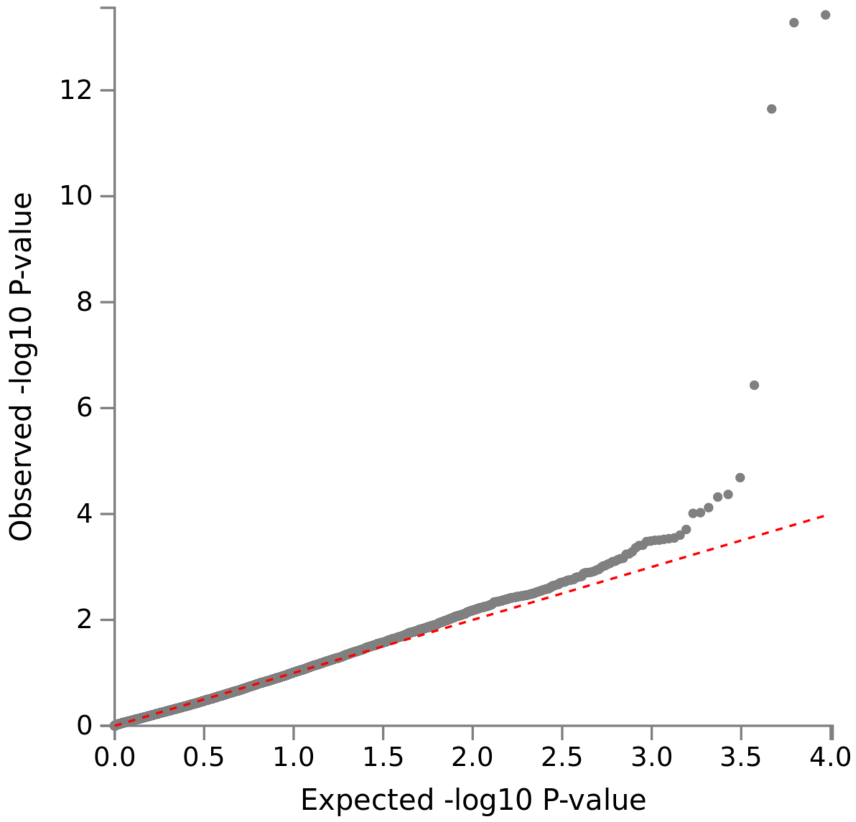

**Figure S5a.** AB_Zscore

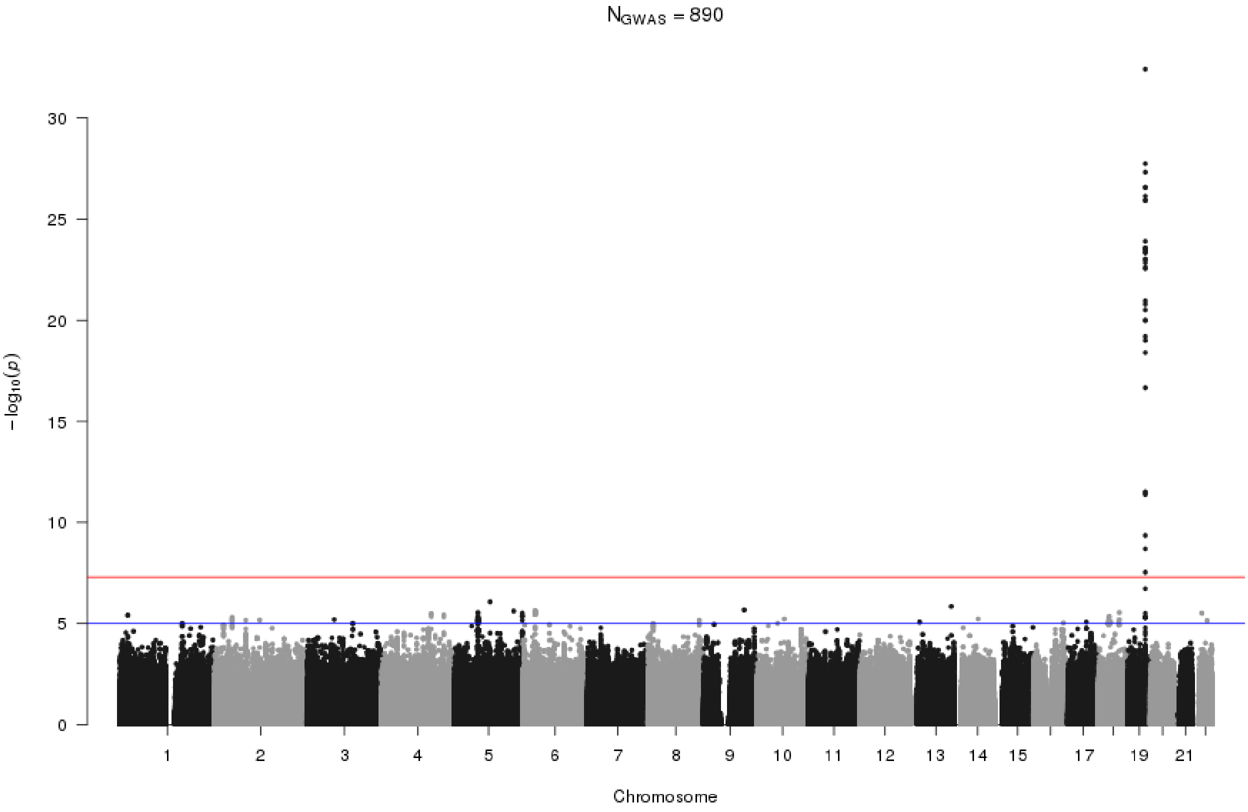


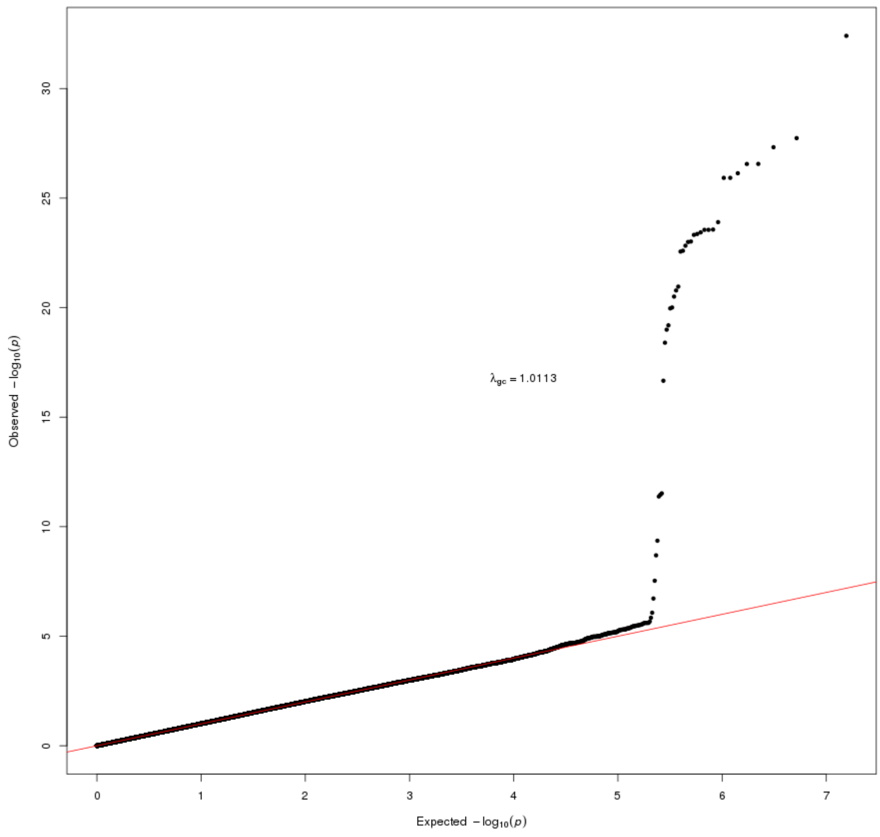

**Figure S5b.** AB_Zscore

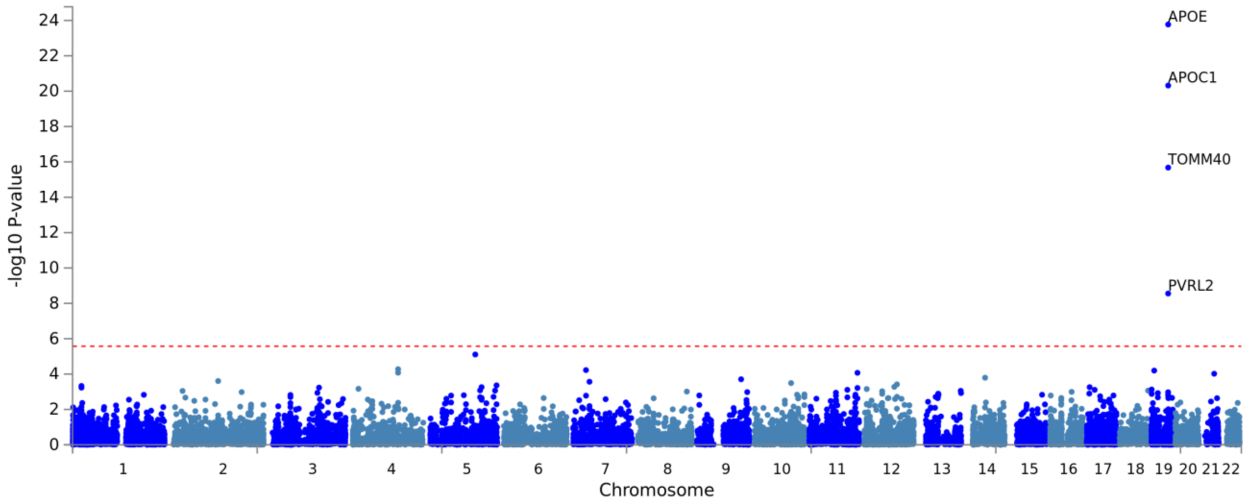


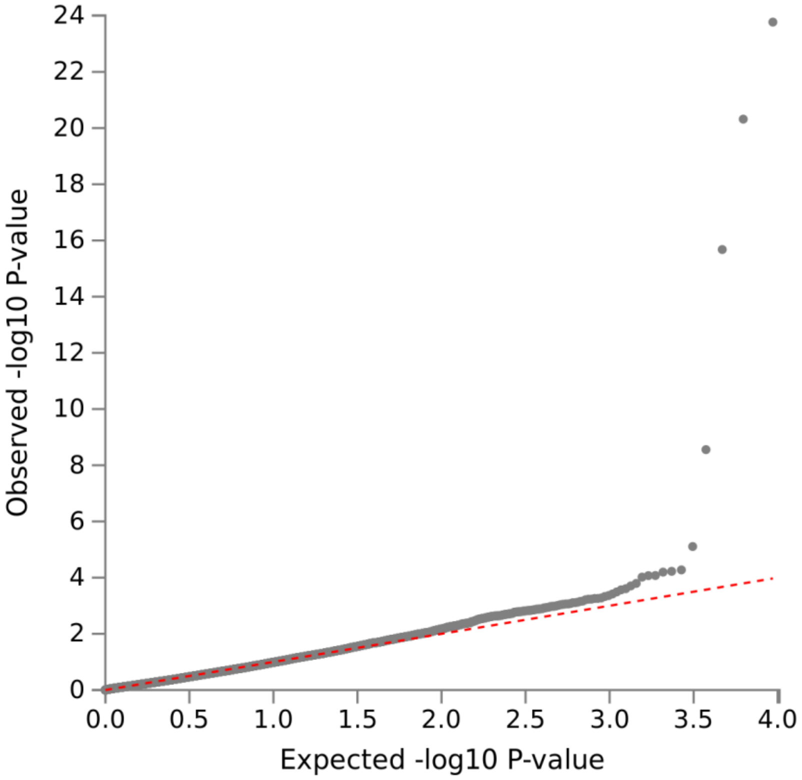

**Figure S6a.** log_Central_CSF_AB42

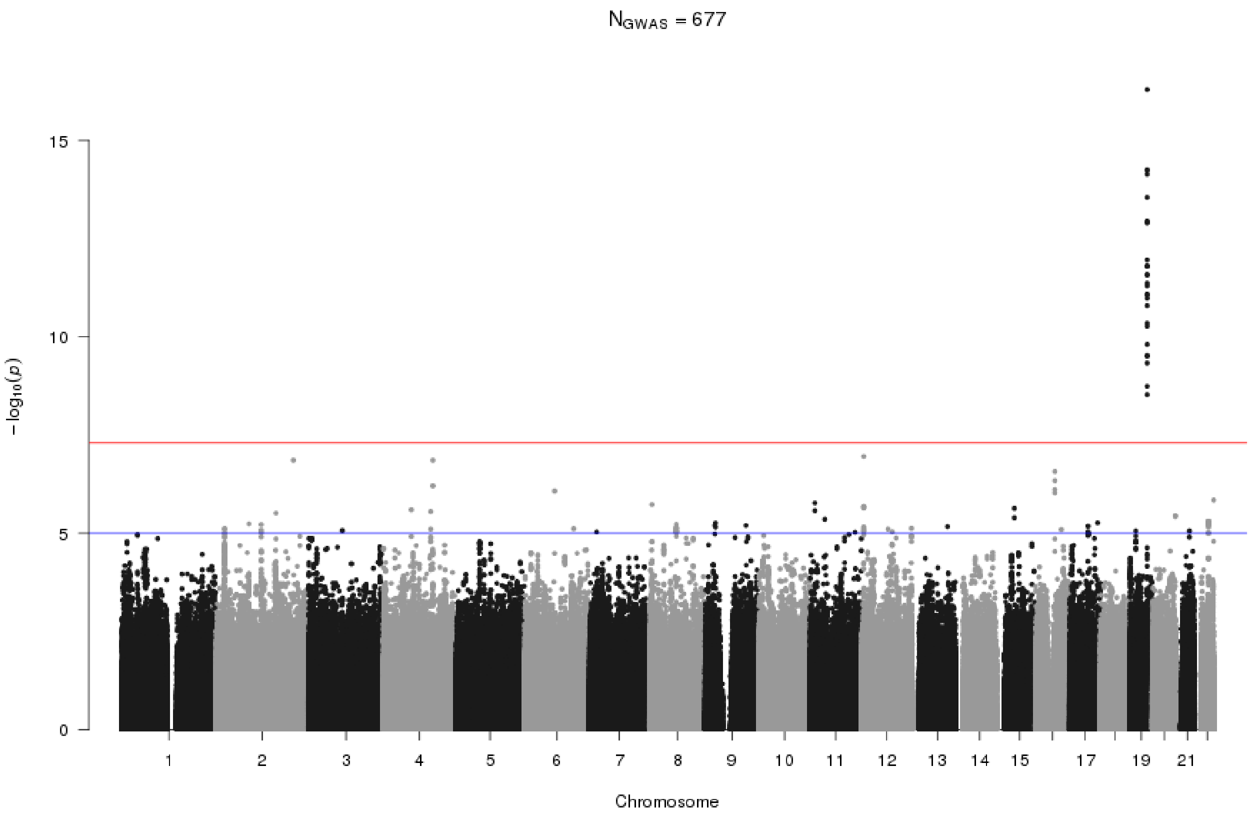


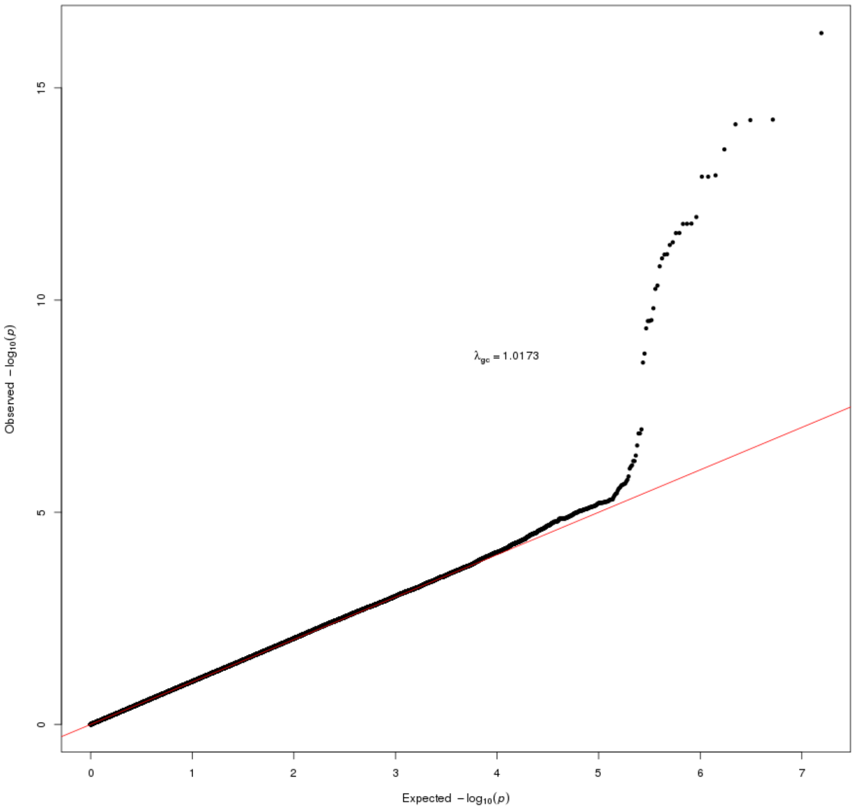

**Figure S6b.** log_Central_CSF_AB42

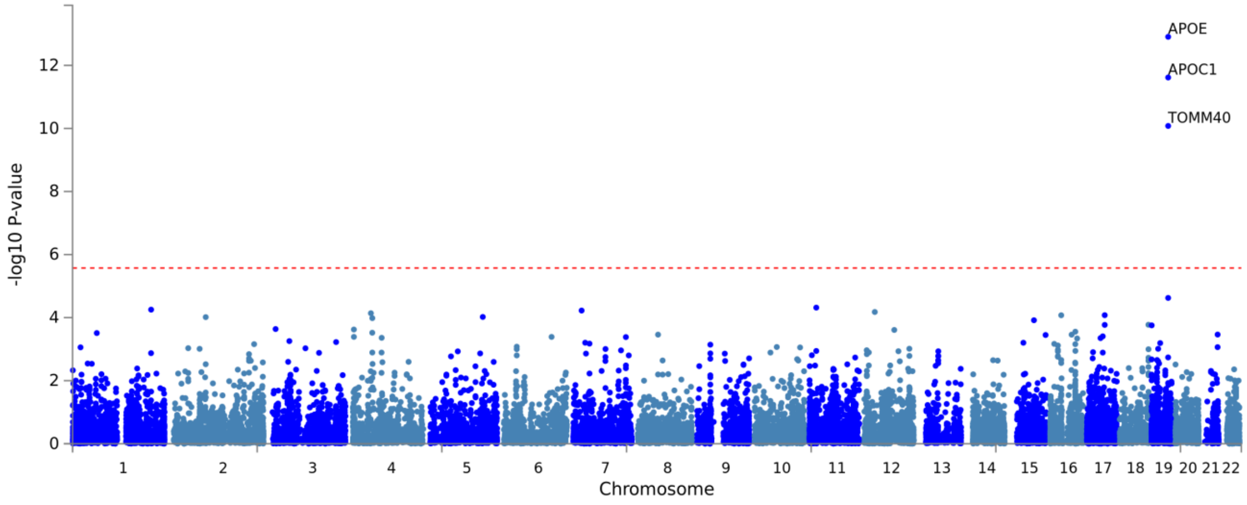


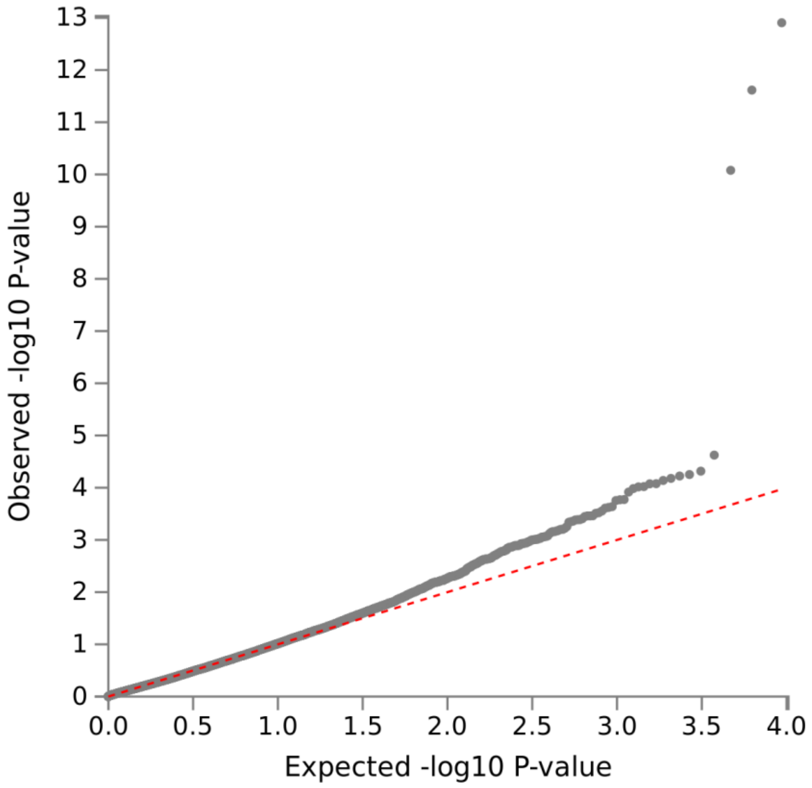

**Figure S7a.** log_Central_CSF_AB4240ratio

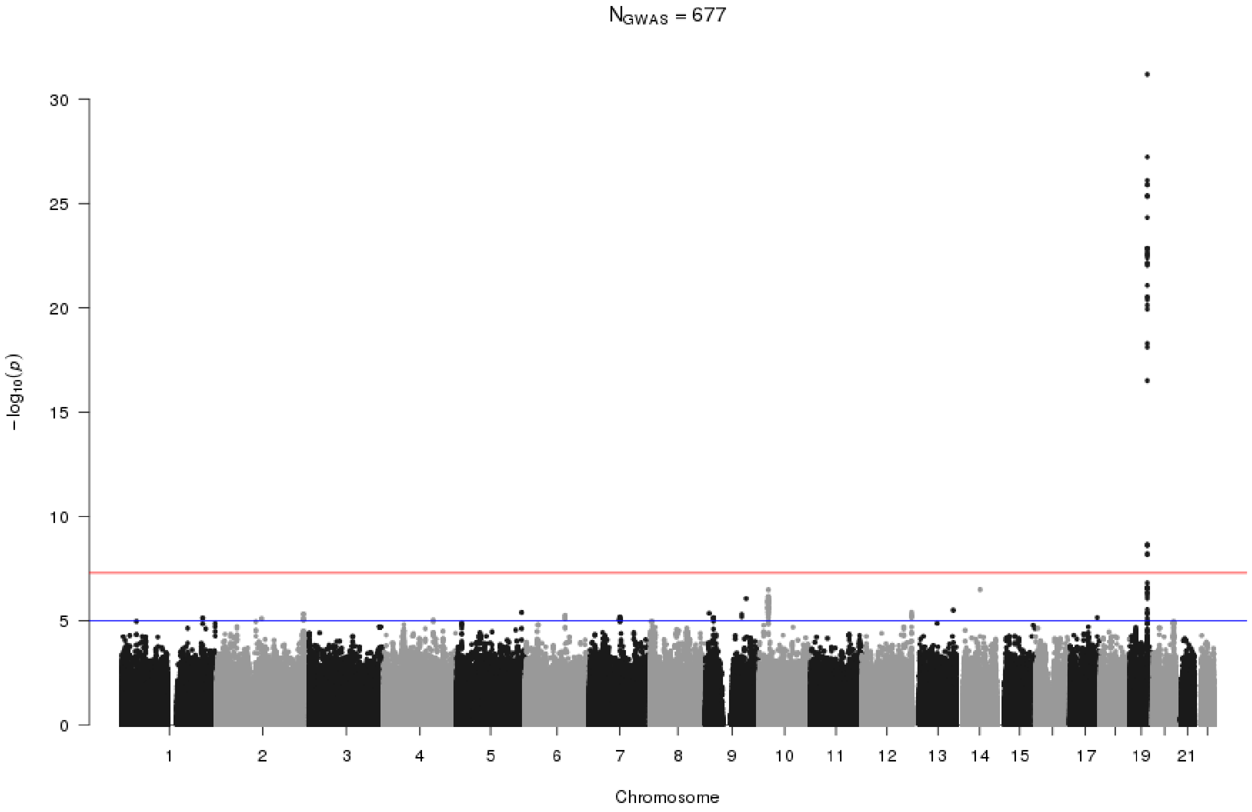


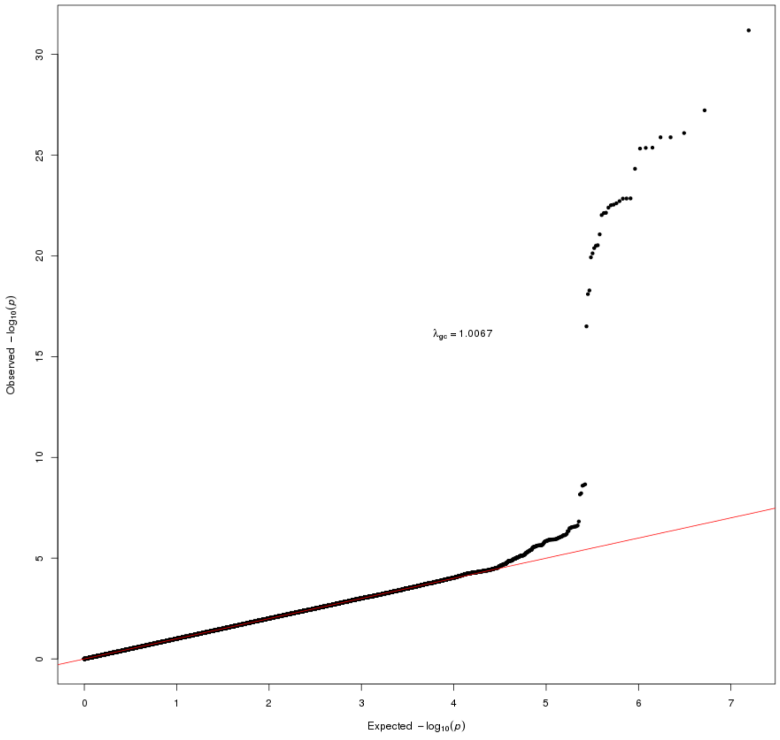

**Figure S7b.** log_Central_CSF_AB4240ratio

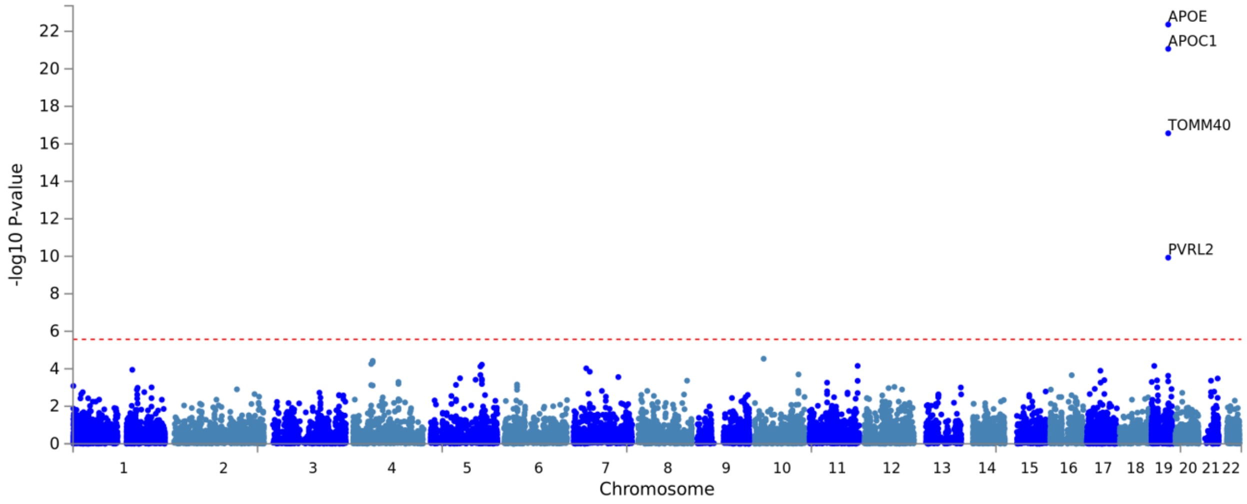


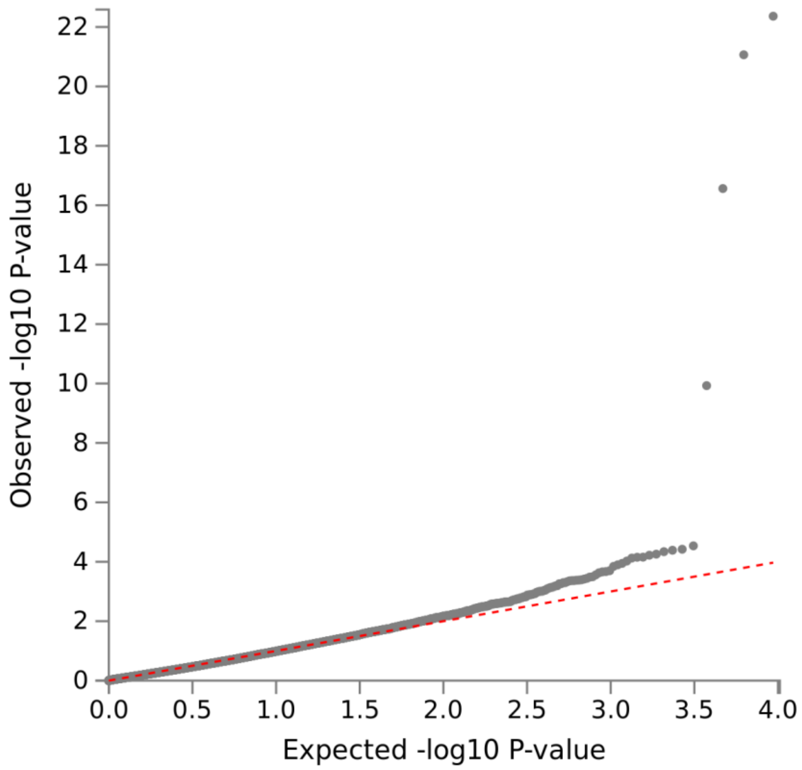

**Figure S8a.** CSF-Ab38

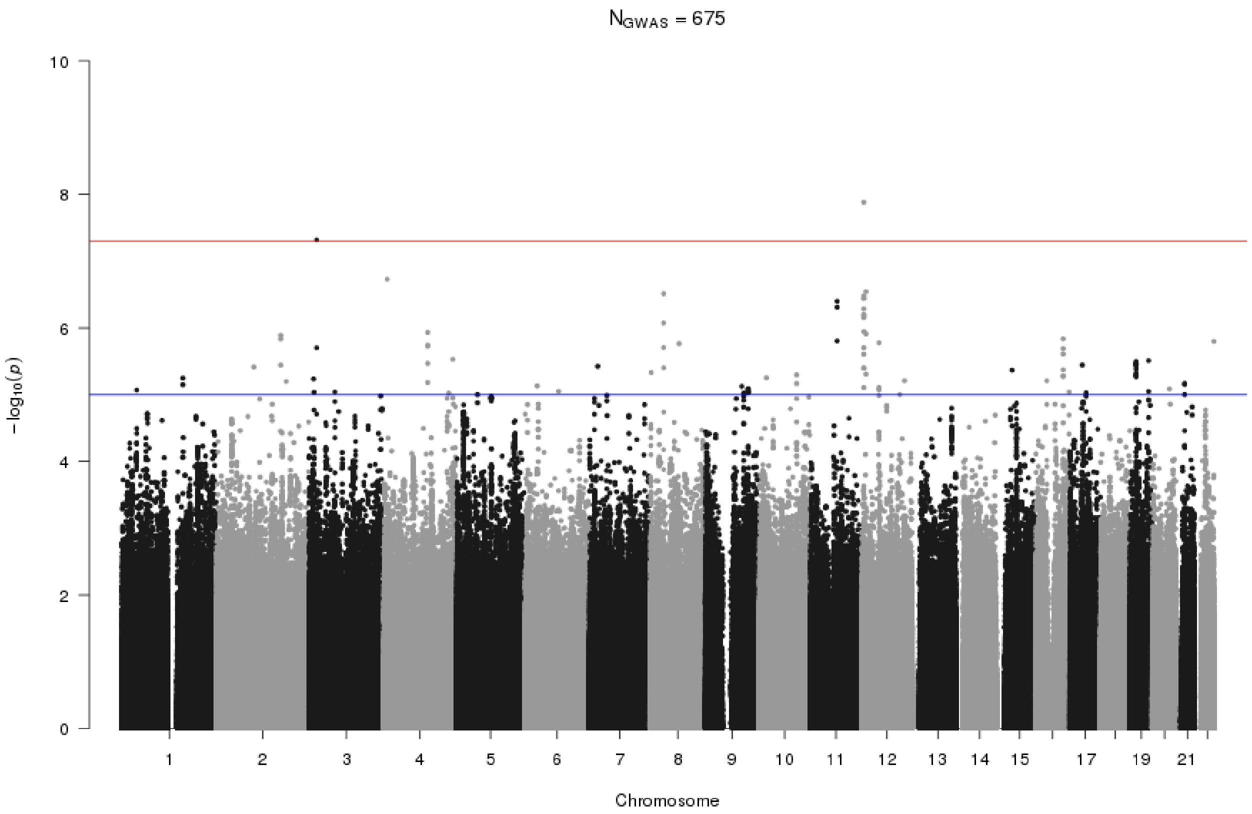


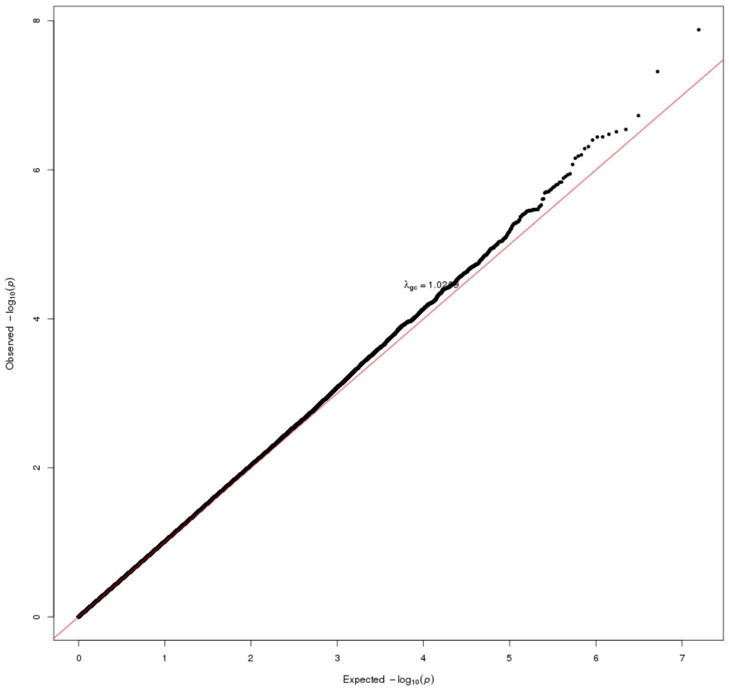

**Figure S8b.** CSF-Ab38

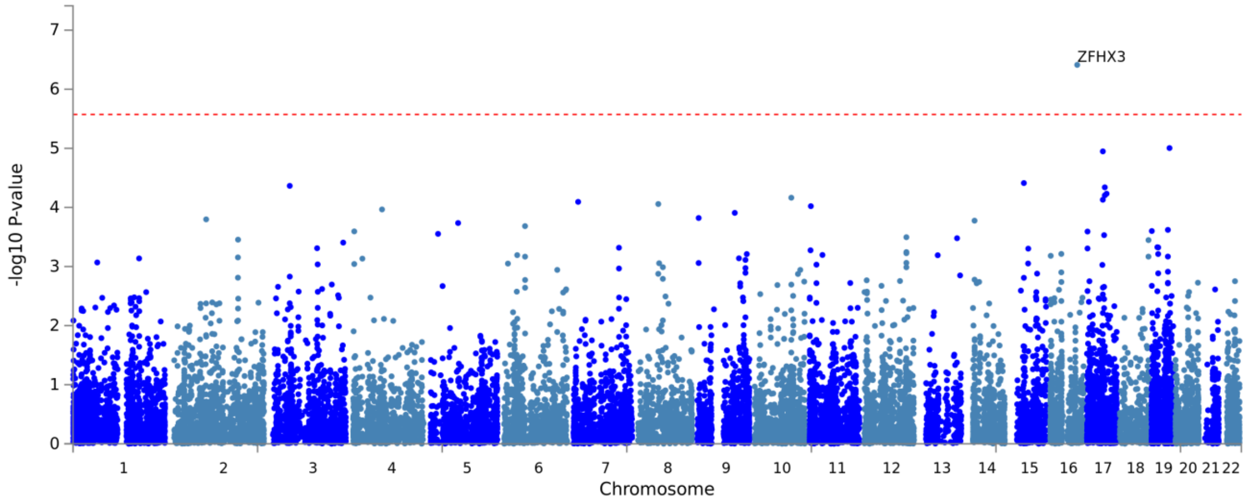


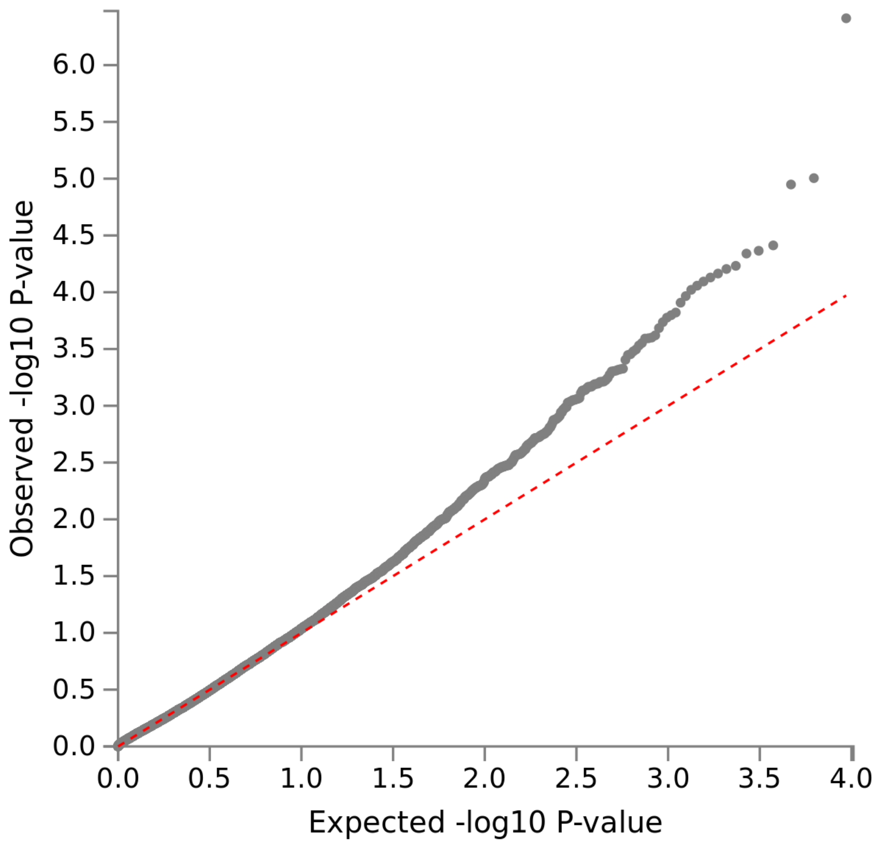

**Figure S9a.** CSF-Ab40

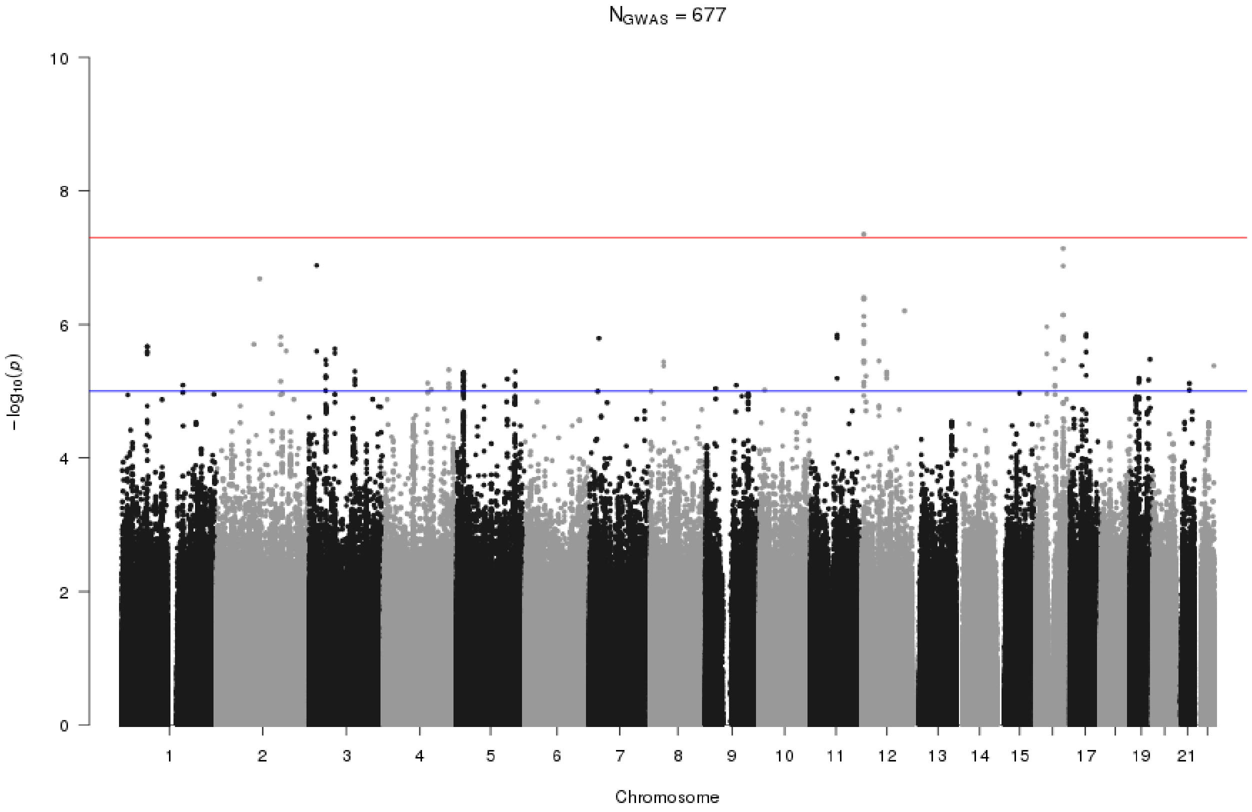


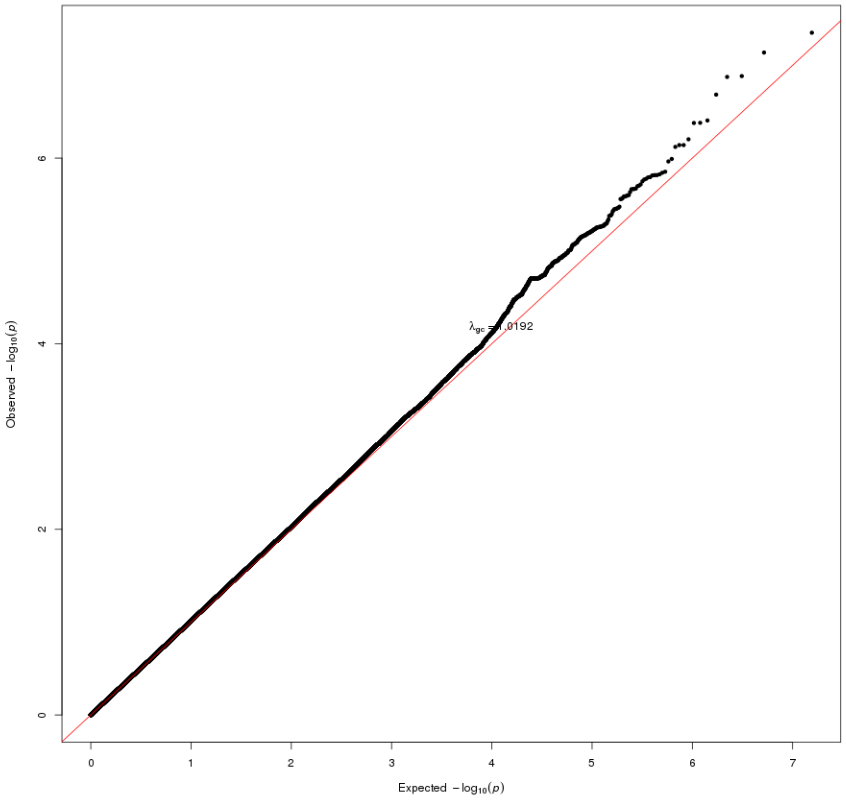

**Figure S9b.** CSF-Ab40

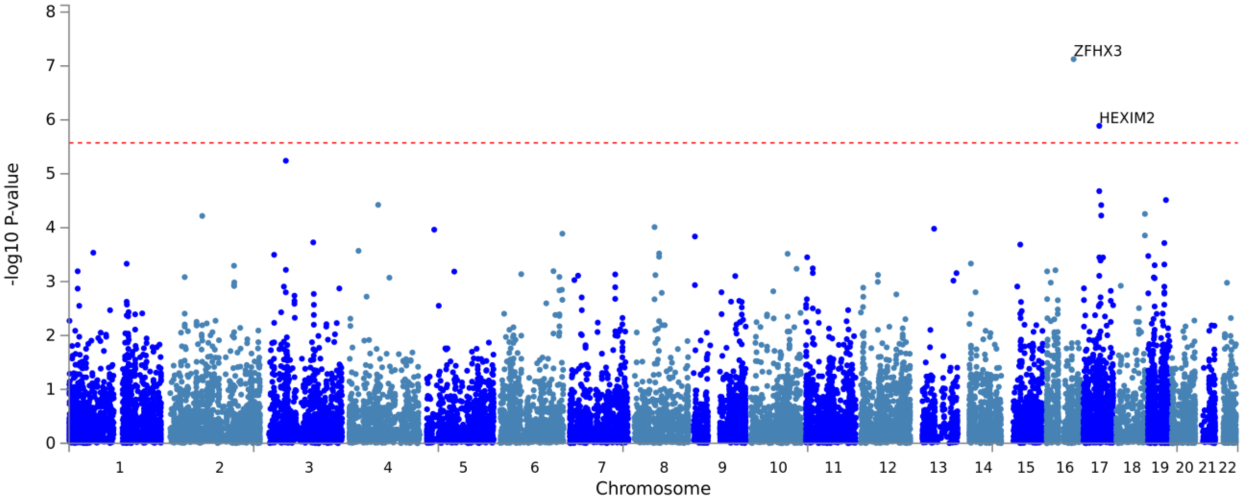


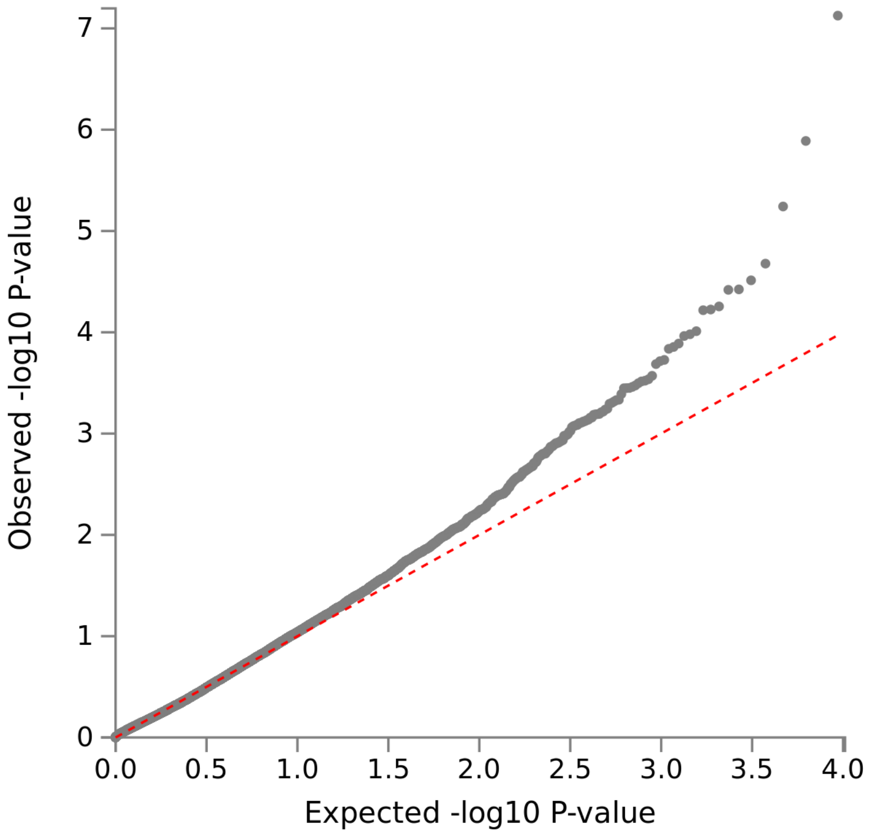

**Figure S10a.** Local_TTAU_Abnormal

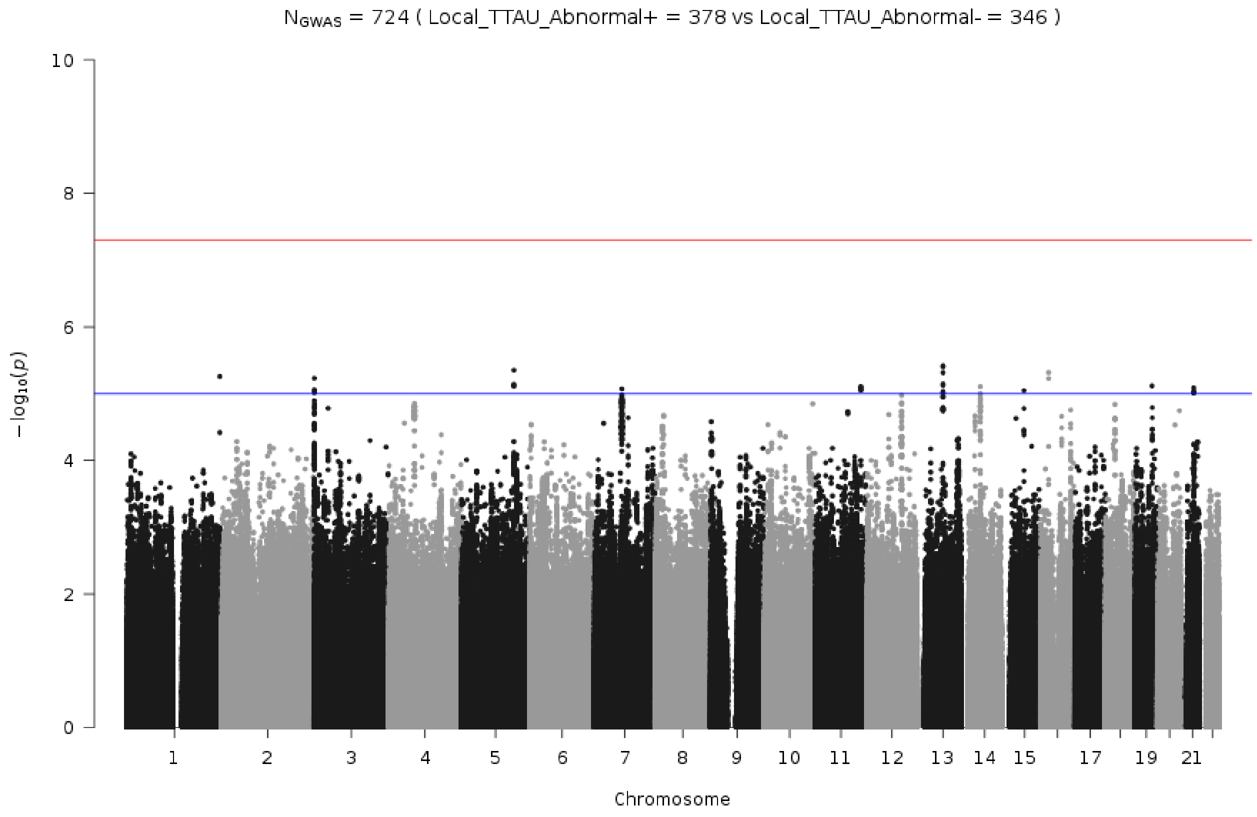


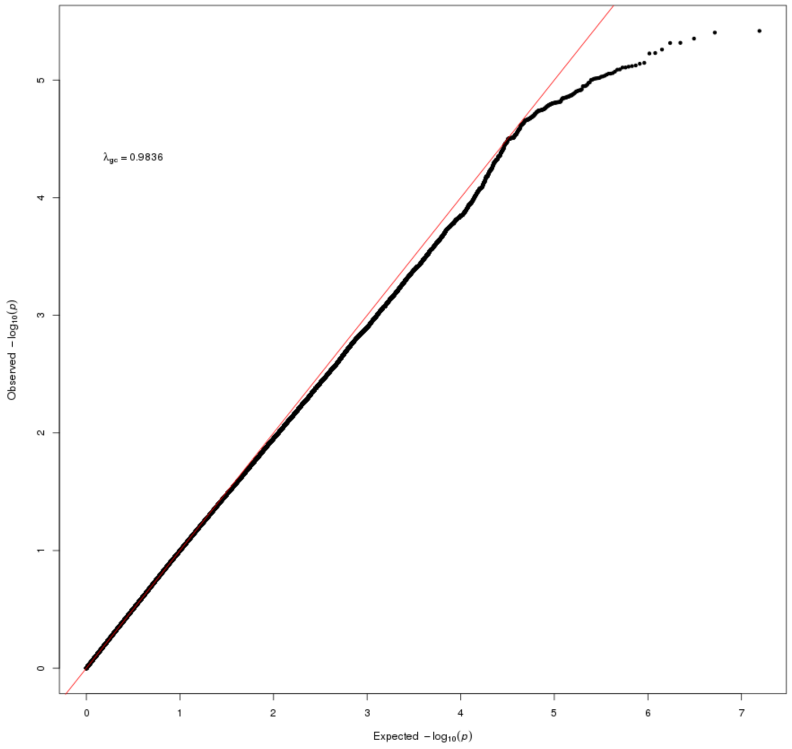

**Figure S10b.** Local_TTAU_Abnormal

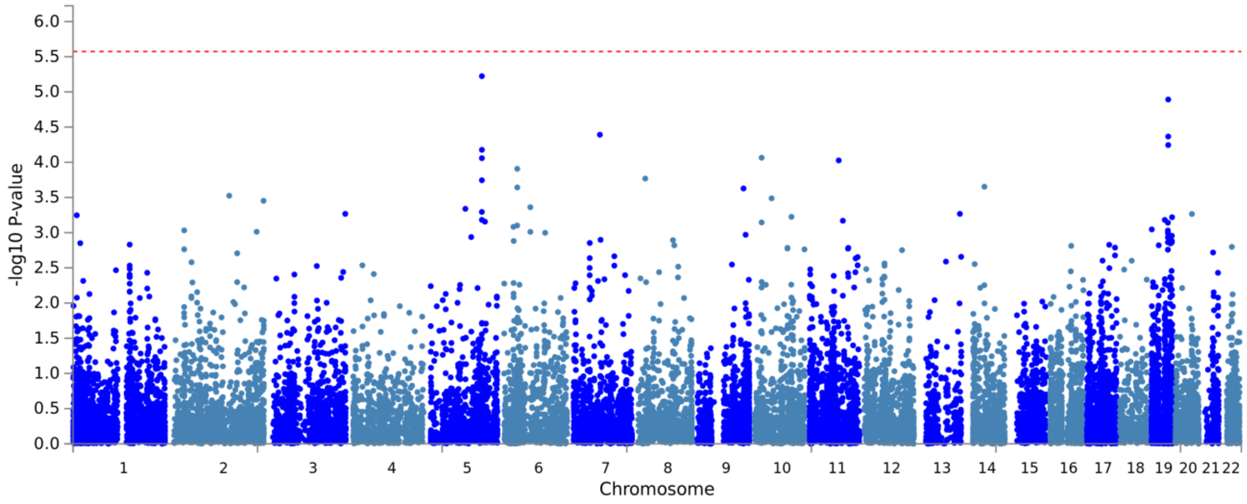


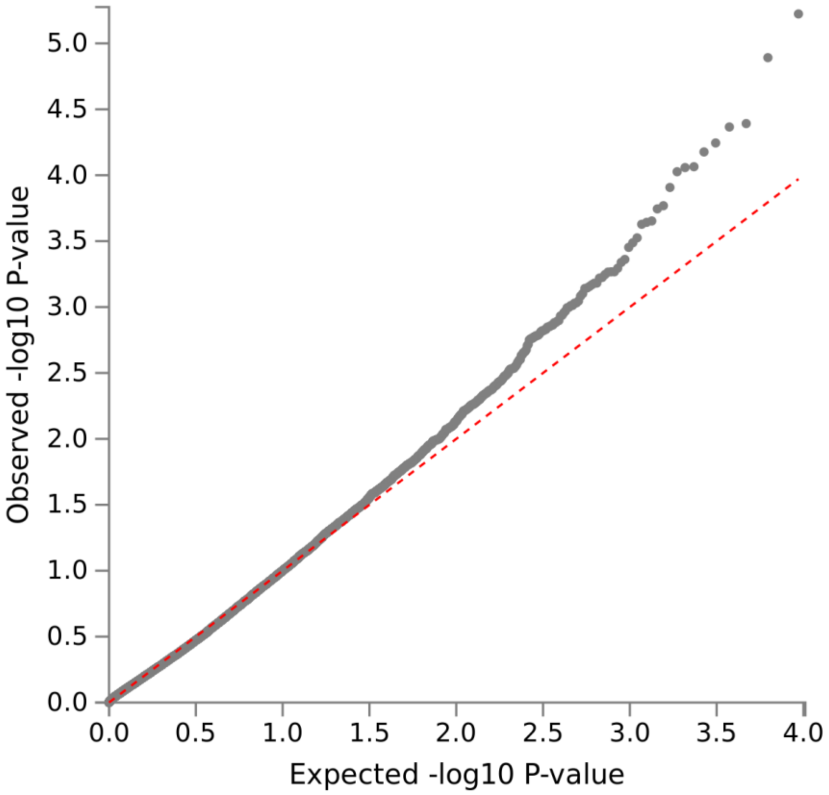

**Figure S11a.** Ttau_ASSAY_Zscore

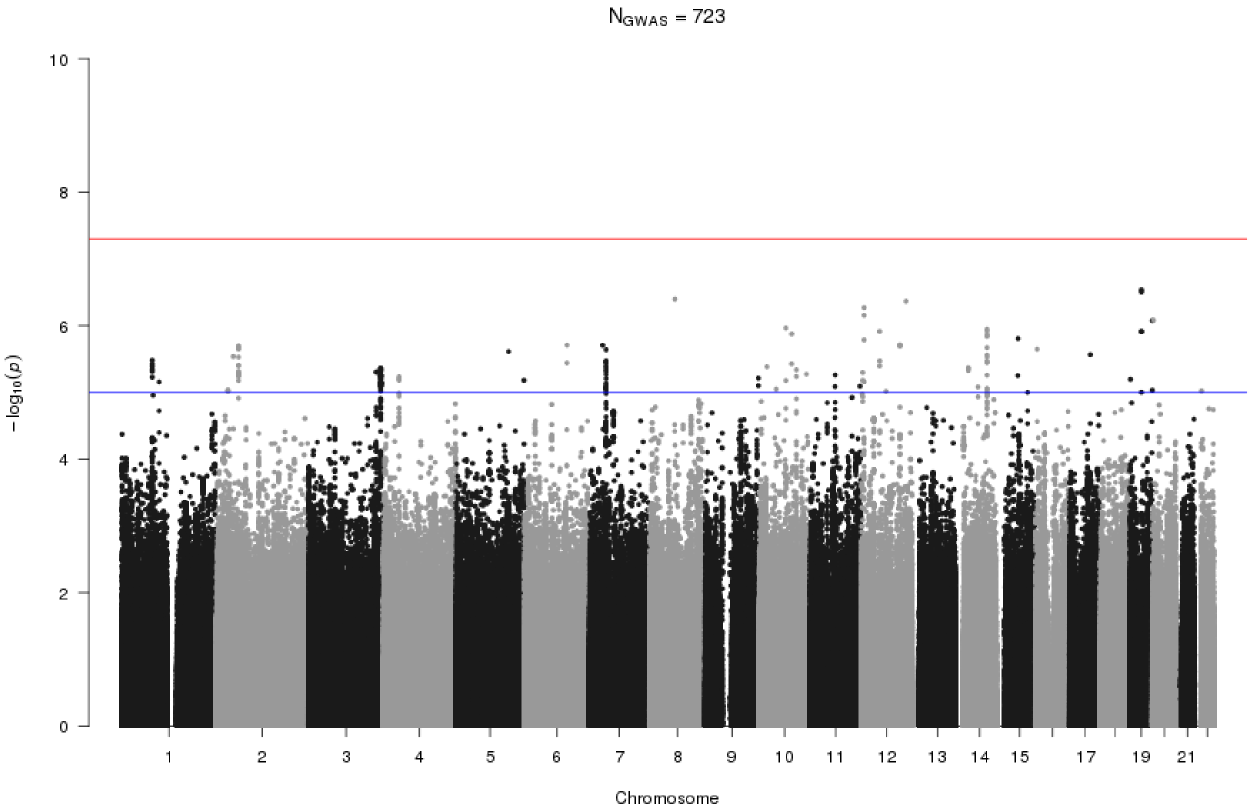


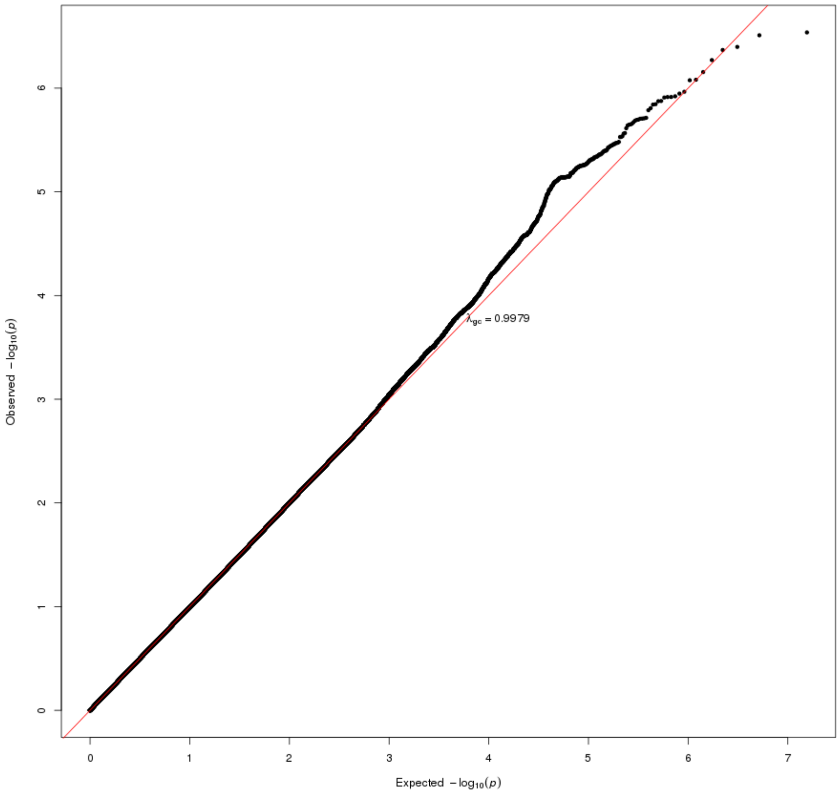

**Figure S11b.** Ttau_ASSAY_Zscore

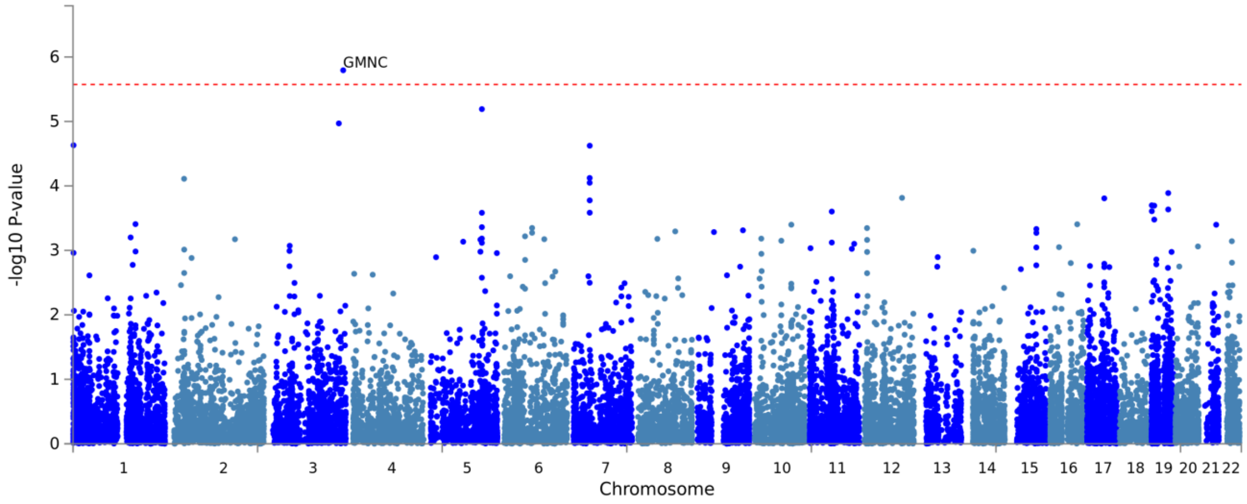


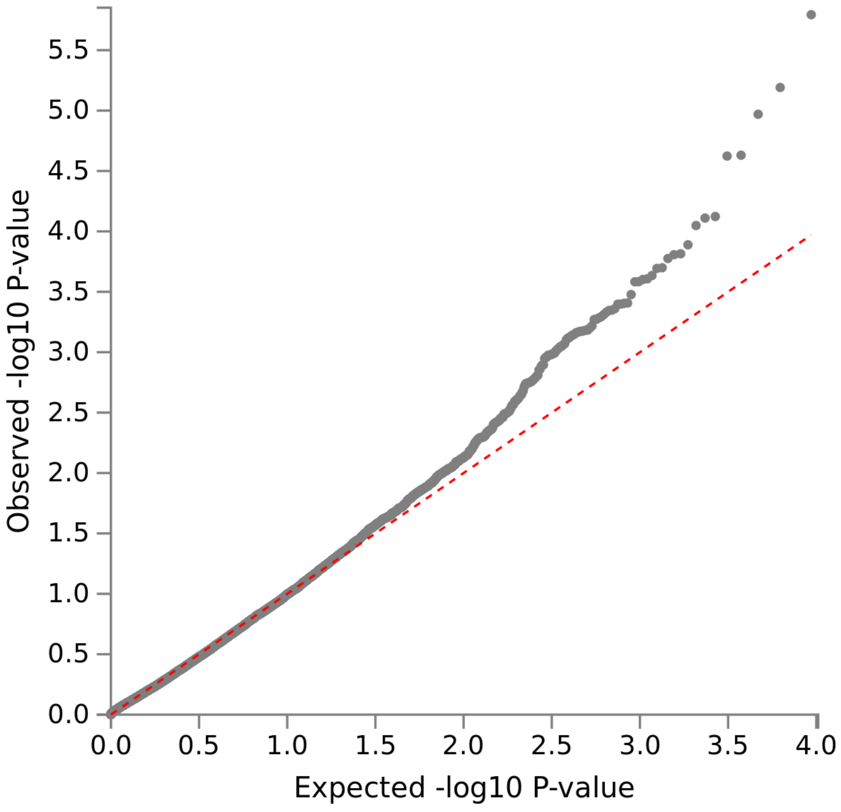

**Figure S12a.** Local_PTAU_Abnormal

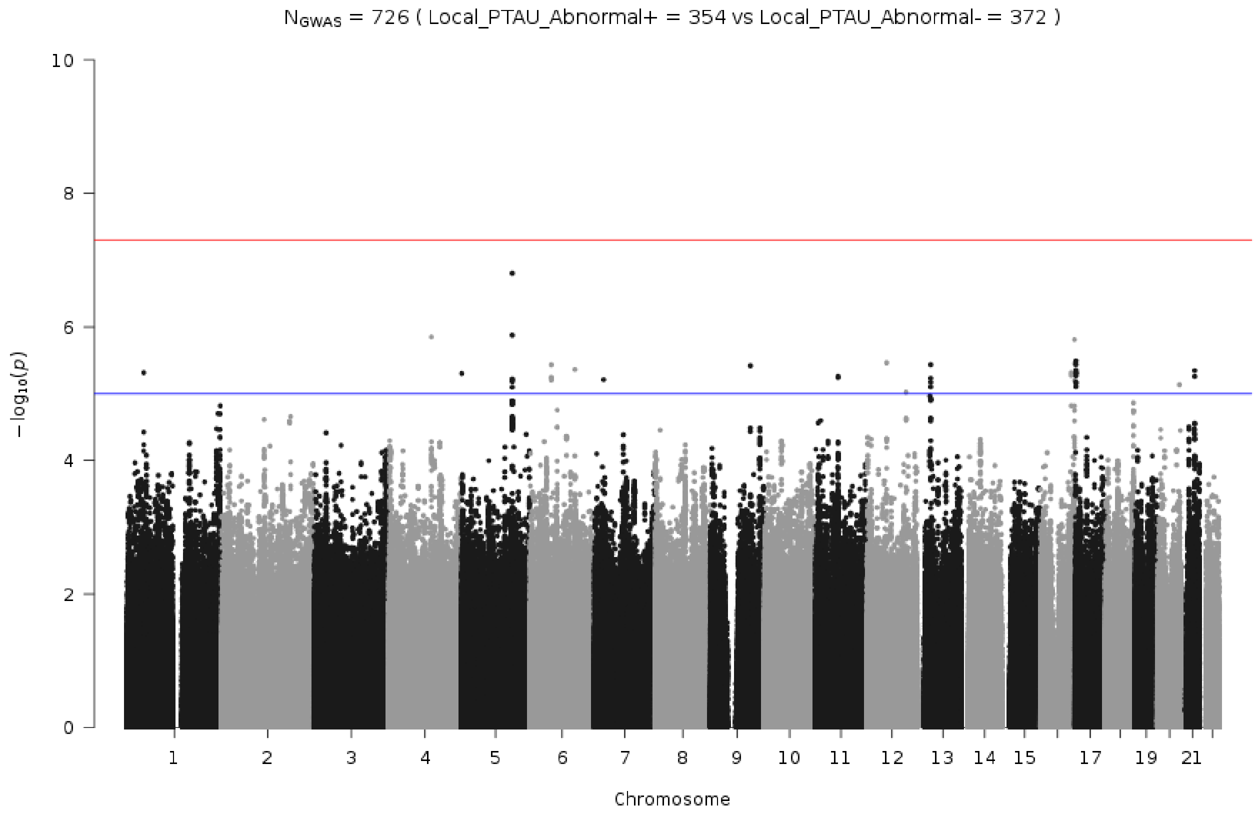


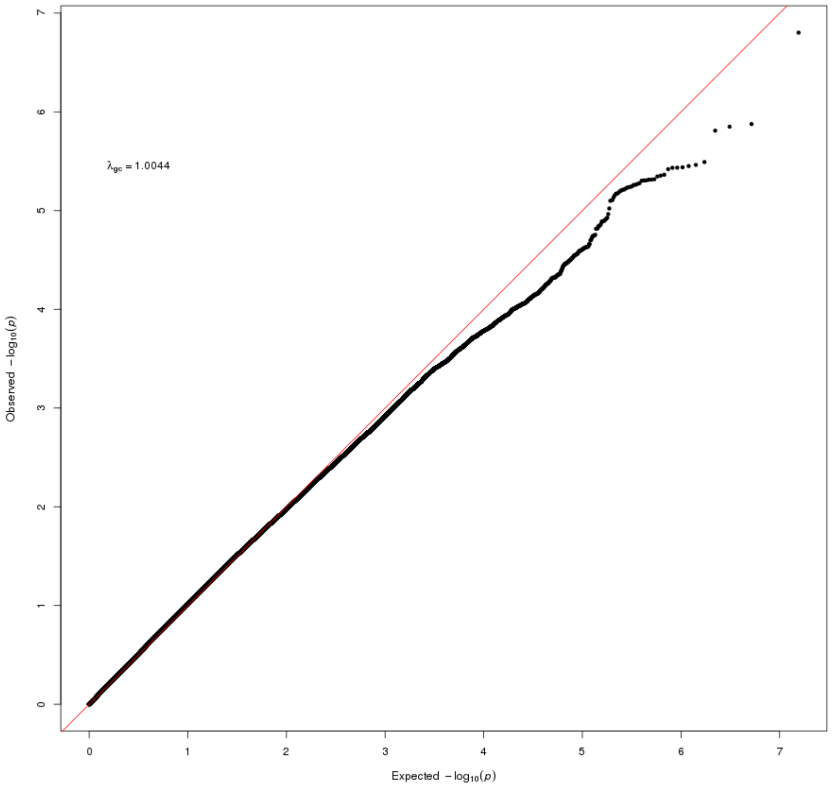

**Figure S12b.** Local_PTAU_Abnormal

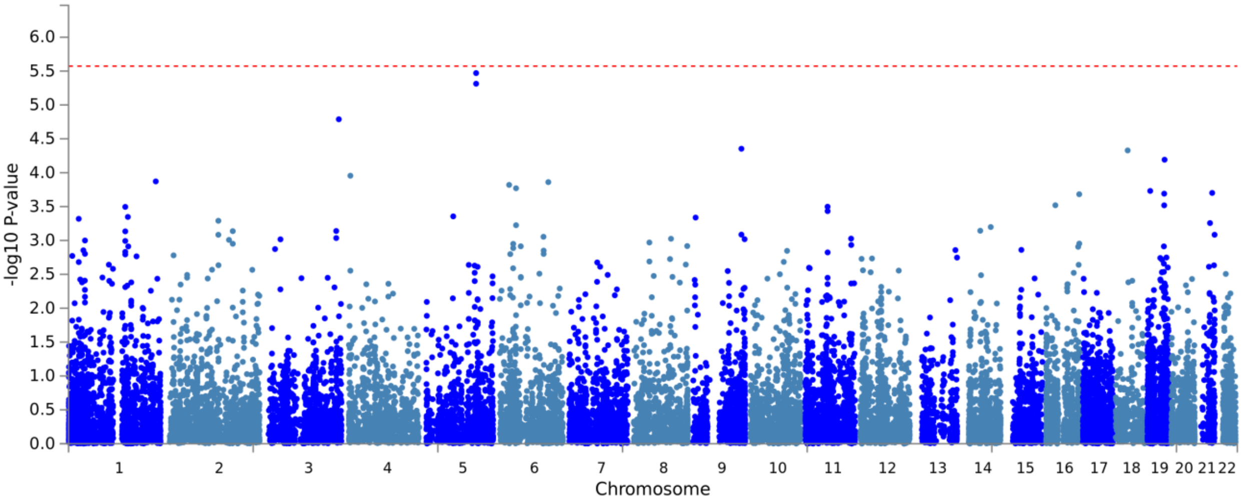


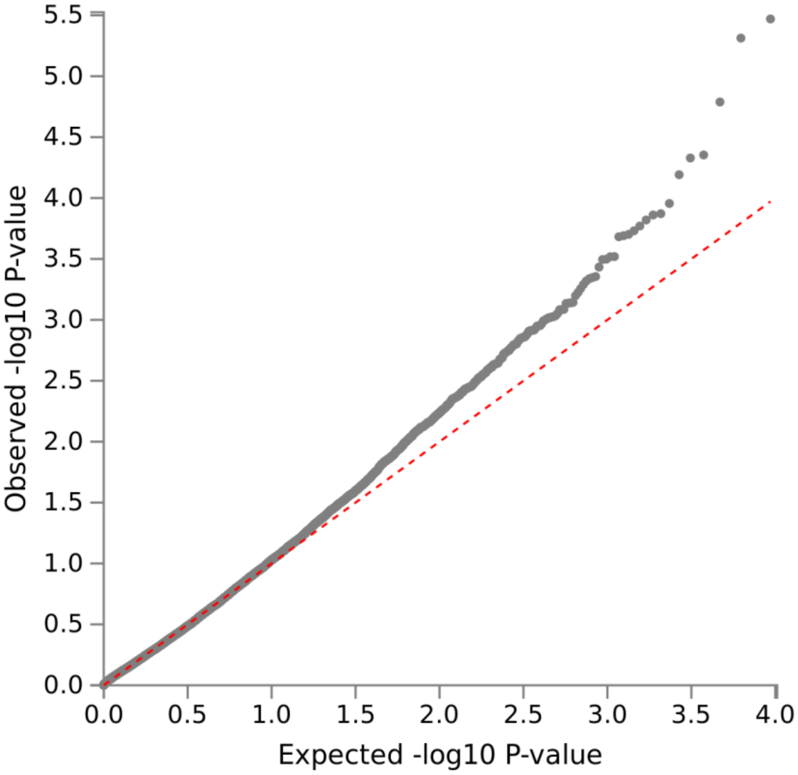

**Figure S13a.** Ptau_ASSAY_Zscore

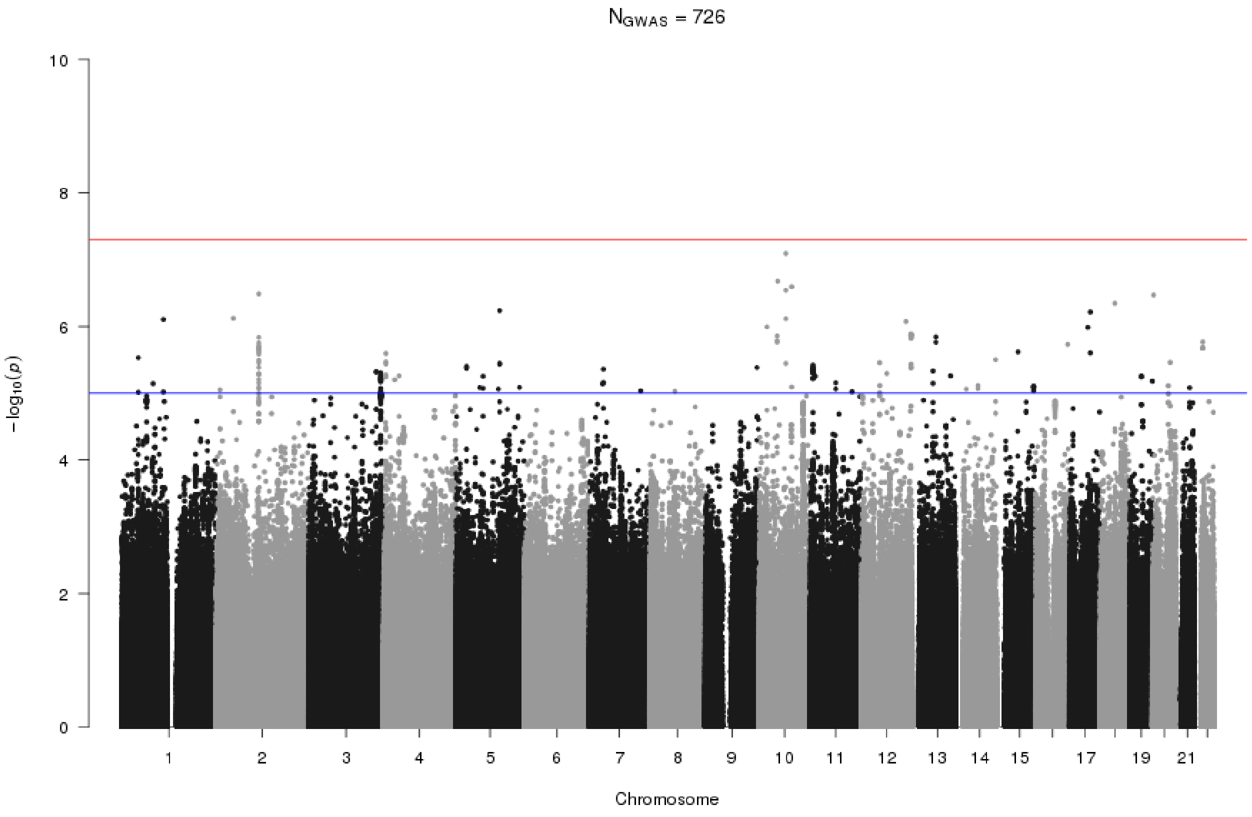


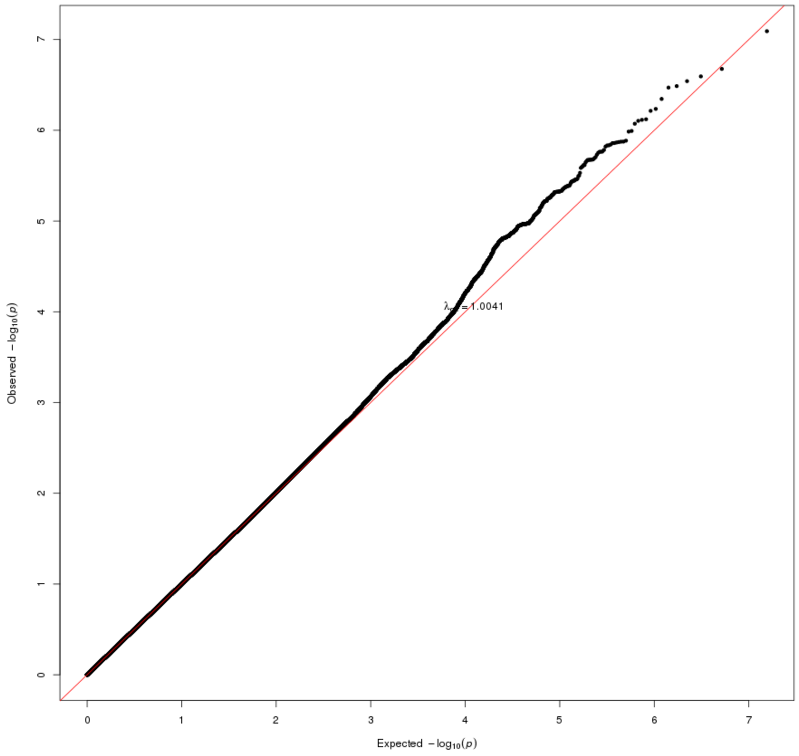

**Figure S13b.** Ptau_ASSAY_Zscore

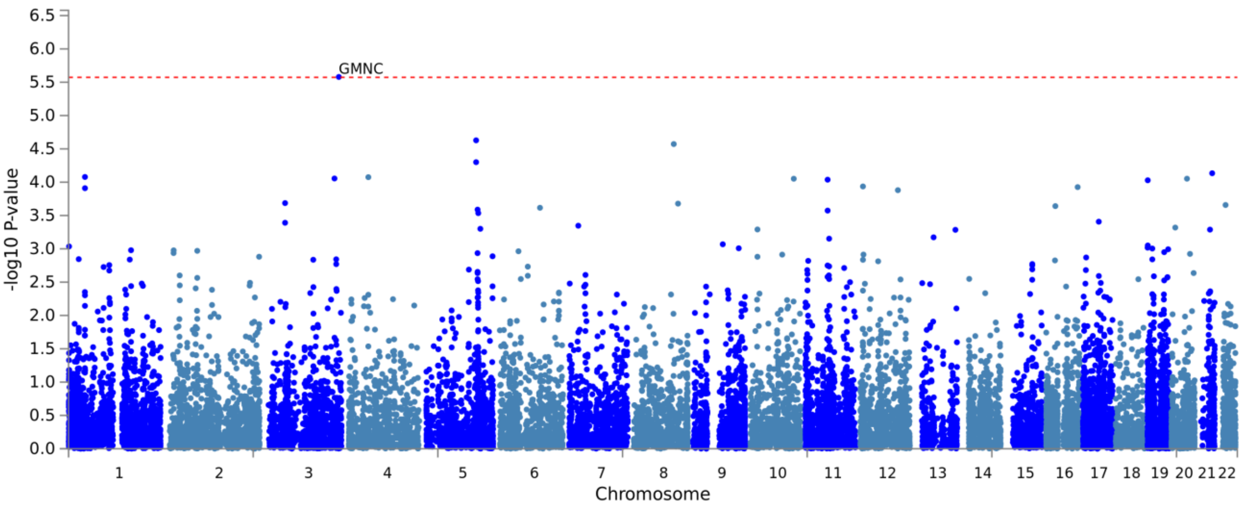


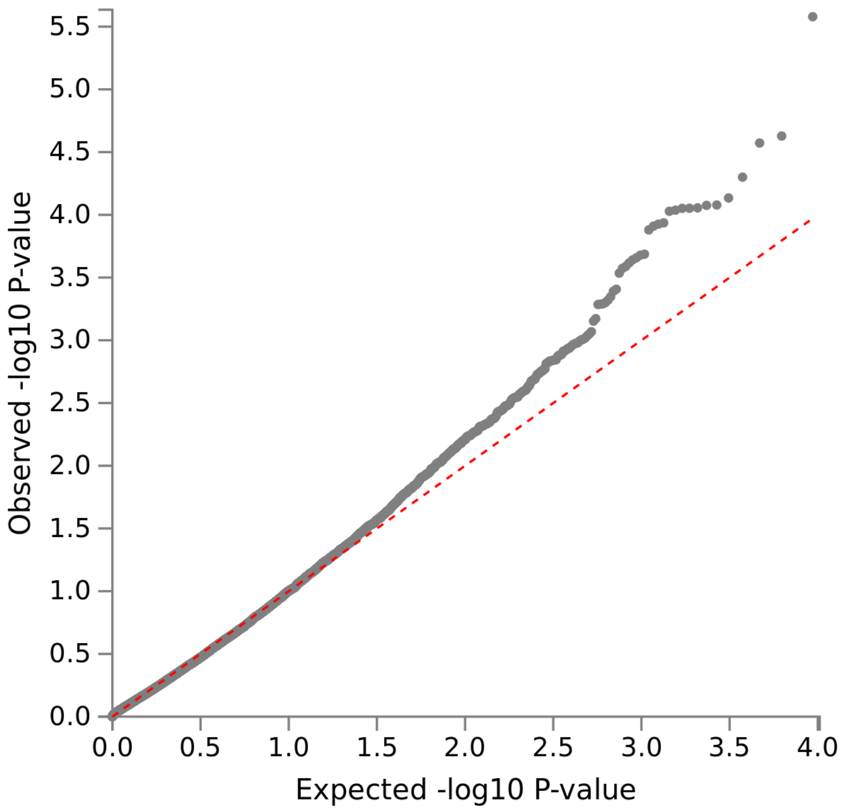
